# HAPSTR1 localizes HUWE1 to the nucleus to limit stress signaling pathways

**DOI:** 10.1101/2023.01.26.525555

**Authors:** Julie K Monda, Xuezhen Ge, Moritz Hunkeler, Katherine A Donovan, Michelle W Ma, Cyrus Y Jin, Marilyn Leonard, Eric S Fischer, Eric J Bennett

## Abstract

HUWE1 is a large, enigmatic HECT domain ubiquitin ligase implicated in the regulation of diverse pathways including DNA repair, apoptosis, and differentiation. How HUWE1 engages its structurally diverse substrates and how HUWE1 activity is regulated are unknown. Using unbiased quantitative proteomics, we identify C16orf72/HAPSTR1 as a dedicated HUWE1 substrate despite HUWE1 targeting substrates in a largely cell type-specific manner. The established physical and genetic interactions between HUWE1 and C16orf72/HAPSTR1 suggest that HAPSTR1 positively regulates HUWE1 function. Here, we show that HAPSTR1 is both a HUWE1 substrate and is required for HUWE1 nuclear localization to facilitate HUWE1 nuclear substrate targeting. HUWE1 or HAPSTR1 loss of function triggers a broad transcriptional stress response. We show that nuclear HUWE1 is both critical for modulating stress signaling pathways, which include p53 and NF-κB-mediated signaling and required for cell proliferation. Combined, our results define a role for HAPSTR1 in gating critical nuclear HUWE1 functions.

## Introduction

Ubiquitin ligases directly engage substrates and provide specificity within the protein ubiquitylation cascade (Yau and Rape, 2016; Zheng and Shabek, 2017). With over 600 putative ubiquitin ligases encoded within the human genome, a diversity of substrate targeting mechanisms have been described (Harper and Schulman, 2021; Wang et al., 2020; Zheng and Shabek, 2017). Substrate ubiquitylation is often tightly regulated to prevent spurious targeting of critical substrates and ensure precise pathway regulation. Some ligases reportedly engage a highly diverse set of substrates with undefined mechanisms (Brunet et al., 2020; Kim et al., 2021; Qi et al., 2022; Wang et al., 2020). How these broadly acting ligases engage their substrates, and how these ligases are regulated remain open questions.

HUWE1 is one such enigmatic ubiquitin ligase that has many reported substrates regulating highly diverse cellular pathways (Giles and Grill, 2020; Qi et al., 2022). HUWE1 is a giant and highly abundant HECT-domain ubiquitin ligase with a variety of protein interaction domains that have been shown to target specific substrates (Adhikary et al., 2005; Chen et al., 2005; Liu et al., 2005; Zhong et al., 2005). Recent structural characterization of full-length human HUWE1 revealed a solenoid architecture built by armadillo repeats to form a central cavity, and the HECT domain positioned above the ring plane (Hunkeler et al., 2021). The characterized HUWE1 substrate, DDIT4 (Thompson et al., 2014), was shown to bind to the inner armadillo repeats within HUWE1 suggesting a new mode of substrate interaction that may be flexible enough to allow for structurally diverse substrate engagement (Hunkeler et al., 2021). This type of plasticity with regards to substrate engagement may be desirable for a subset of ligases, like quality control ligases that need to target a diverse range of damaged or misfolded substrates. Indeed, HUWE1 has been shown to act in a quality control manner to target orphan proteins of multi-subunit complexes, such as unassembled ribosomal proteins and histone proteins, for degradation (Defenouillere et al., 2017; Grabarczyk et al., 2021; Singh et al., 2012; Sung et al., 2016). However, how HUWE1 and similar ubiquitin ligases with demonstrated substrate plasticity are regulated to prevent spurious substrate degradation is incompletely understood.

One mechanism that is employed to regulate ubiquitin ligase function is to restrict ligase subcellular localization. This feature is characteristic of ligases that regulate endoplasmic reticulum (ER) function (Sun and Brodsky, 2019). HUWE1 is broadly localized to both the cytoplasm and the nucleoplasm and has many reported substrates within each compartment (Atsumi et al., 2015; Cassidy et al., 2020; Chen et al., 2005; Choe et al., 2016; Dominguez-Brauer et al., 2016; Forget et al., 2014; Inoue et al., 2013; Mandemaker et al., 2017; Parsons et al., 2009; Thompson et al., 2014; Wang et al., 2014; Zhao et al., 2009; Zhao et al., 2008; Zhong et al., 2005). HUWE1 has been shown to regulate many nuclear processes including transcription, DNA damage responses, and stress signaling. HUWE1 can ubiquitylate transcriptional regulators, like c-Myc, N-Myc, and p53 to govern developmental and stress-dependent signaling pathways (Adhikary et al., 2005; Chen et al., 2005; Endres et al., 2021; Hao et al., 2012; King et al., 2016; Kon et al., 2012; Myant et al., 2017; Peter et al., 2014; Yang et al., 2018; Zhao et al., 2009; Zhao et al., 2008). Further, HUWE1 regulates DNA damage responses by ubiquitylating diverse substrates ranging from histones to the DNA repair polymerase λ (Choe et al., 2016; Kunz et al., 2020; Mandemaker et al., 2017; Markkanen et al., 2012; Myant et al., 2017; Parsons et al., 2009). Human mutations within HUWE1 have been linked to intellectual disability disorders and neurodevelopmental defects underscoring the broad role HUWE1 plays in regulating nuclear functions (Froyen et al., 2012; Froyen et al., 2008; Giles and Grill, 2020; Madrigal et al., 2007).

To understand how HUWE1 can target diverse substrates in a controlled manner, we used an unbiased proteomic approach to systematically identify HUWE1 substrates. Confirming the plasticity of HUWE1 targeting, we identify a broad range of new HUWE1 substrates, nearly all of them uncharacterized. Nuclear localized proteins are enriched among the identified HUWE1 substrates, and we identify C16orf72 (also known as TAPR1 and HAPSTR1) as a robust HUWE1 nuclear substrate. Despite being a substrate, C16orf72/HAPSTR1 has a strong positive genetic relationship with HUWE1, in contrast to what is expected for a ubiquitin ligase-substrate relationship. We find that C16orf72/HAPSTR1 is required for HUWE1 nuclear localization and targeting of nuclear substrates. Mutations that disrupt C16orf72/HAPSTR1 nuclear localization or HUWE1 binding compromise HUWE1 nuclear activity. Nuclear HUWE1 impacts stress signaling pathways including NF-κB-mediated inflammatory signaling and p53 signaling. Loss of C16orf72/HAPSTR1 or HUWE1 activates p53 signaling in a manner that depends upon the ability of C16orf72/HAPSTR1 to localize HUWE1 to the nucleus. Further, loss of nuclear HUWE1 activity limits cellular growth, despite total HUWE1 cellular levels remaining unchanged. Together, we demonstrate that critical nuclear HUWE1 ubiquitylation activity is enabled by C16orf72/HAPSTR1.

## Results

### Proteomic profiling of HUWE1 substrates

HUWE1 has been reported to regulate a diverse and growing number of cellular pathways through the targeted degradation of structurally diverse proteins (Giles and Grill, 2020; Qi et al., 2022). In order to characterize the mechanisms HUWE1 utilizes to enable such plasticity towards its substrates, we performed quantitative and deep whole cell proteomics to identify proteins whose steady-state abundance is elevated upon HUWE1 loss of function. We generated 293T cells with stable inducible expression of shRNAs targeting HUWE1 and validated that induced expression of the shRNAs substantially reduced HUWE1 protein levels (Figure S1A). We chose the two cell lines with the greatest HUWE1 depletion (155 and 969, hereafter designated shHUWE1-2 and shHUWE1-3, respectively) to perform proteomic profiling of whole cell extracts using a tandem mass tag (TMT) approach. In total, we quantified more than 8800 proteins in both HUWE1 knockdown cell lines and determined protein abundance changes upon HUWE1 knockdown compared to cells expressing an shRNA targeting firefly luciferase (Figures 1A and 1B; Table S1). Only 138 proteins displayed altered abundance (log_2_ fold change greater or less than 0.7) in either cell line, and 104 of the 107 proteins that increased in abundance did not have increased mRNA abundance, indicating that HUWE1 regulates the turnover of these proteins (Table S2). We observed a high correlation between the two HUWE1 shRNA experiments (Figure 1B). Surprisingly, only two previously characterized HUWE1 substrates were identified in this experiment and there was little to no overlap with the previous proteomic experiments using HUWE1 genetic depletion or HUWE1 targeting small molecules to identify HUWE1 substrates (Endres et al., 2021; Thompson et al., 2014). It is possible that the HUWE1 substrate pool is highly variable across cell lines and conditions, or that our HUWE1 depletion, which was around 4-fold, was insufficient to observe significant changes in substrate levels. To directly address the second possibility, we obtained two previously generated and characterized HUWE1 knockout cell lines generated in HAP1 cells (Hundley et al., 2021). We again determined changes in protein abundance comparing HAP1 parental and HUWE1 KO cells using deep quantitative proteomics. From more than 7300 proteins quantified, we identified 227 proteins that increased in abundance (log_2_ fold change greater than 0.7) in one of the two HUWE1 knockout clones compared to parental HAP1 cells (Figure 1C; Table S3). A correlation between the two HUWE1 knockout clones was observed, although to a reduced extent than observed using shRNA mediated HUWE1 depletion (Figure 1C). Despite the observation that HUWE1 protein levels were depleted 16-fold (limit of detection), few known HUWE1 substrates were identified. Further, although 71% of proteins identified from the HUWE1 knockdown experiment in 293T cells were also identified in HAP1 cells, only 21% (23/107) of the proteins that increased in abundance upon HUWE1 shRNA-mediated knockdown in this study also had elevated protein levels in HUWE1 knockout HAP1 cells. Taken together, our data suggests that the HUWE1 substrate pool varies substantially between cell types and likely varies based on cellular conditions.

**Figure 1.**
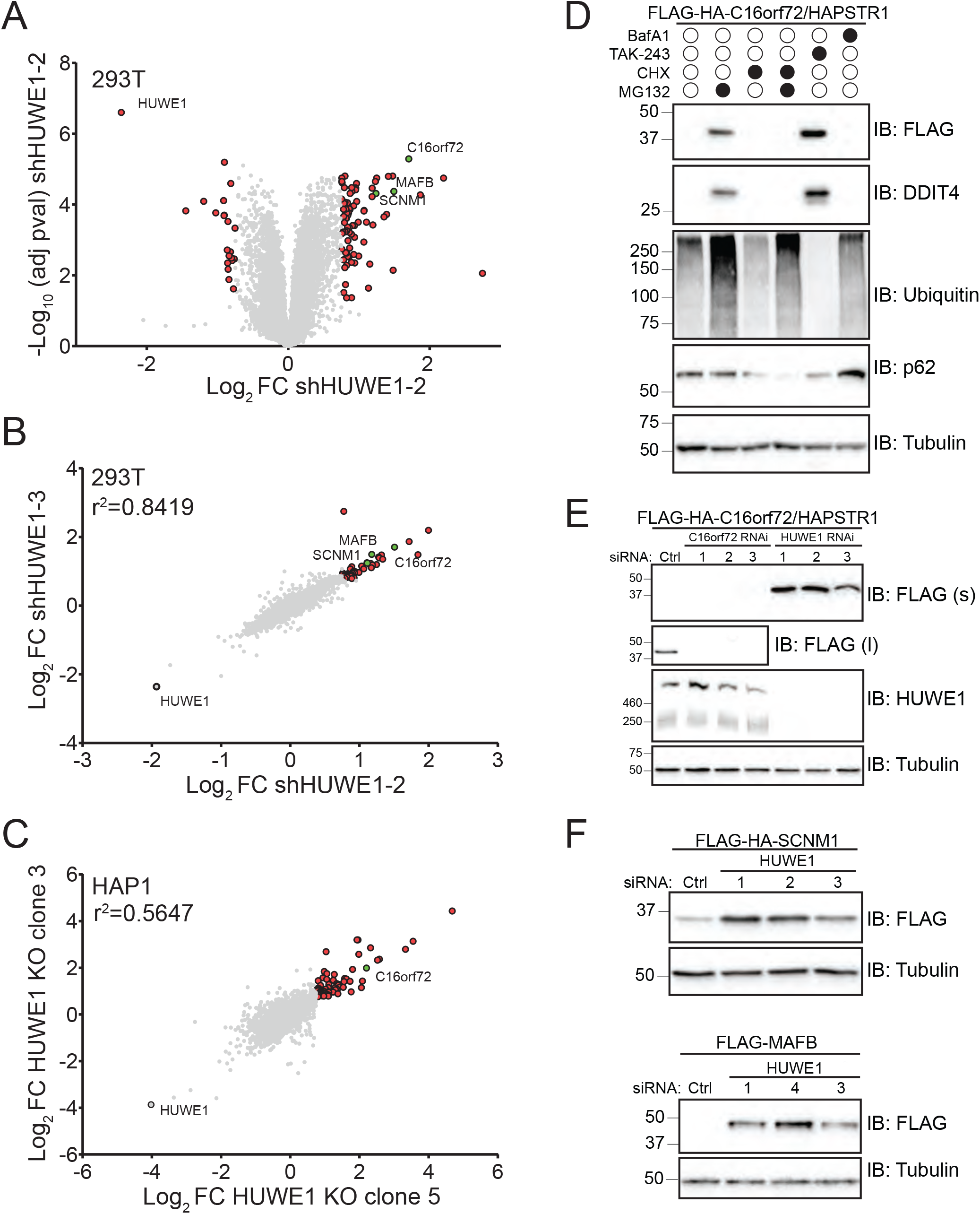
Identification of HUWE1 substrates using quantitative proteomics. A) Volcano plot depicting the log_2_ fold change (FC) and -log_10_ P-value for proteins quantified by tandem mass tag proteomics in 293T cells after 96 hours of doxycycline-inducible expression of an shRNA targeting HUWE1 relative to a firefly luciferase knockdown. Proteins with altered abundance in response to HUWE1 knockdown are indicated in red. Proteins selected for further analysis are indicated in green. B) Scatter plot comparing the log_2_ fold change (FC) for all proteins quantified by tandem mass tag proteomics in 293T cells after 96 hours of doxycycline-inducible expression of two different shRNAs targeting HUWE1 relative to a firefly luciferase knockdown. Proteins with altered abundance in response to HUWE1 knockdown are indicated in red. Proteins selected for further analysis are indicated in green. C) Scatter plot comparing the log_2_ fold change (FC) for all proteins quantified by tandem mass tag proteomics in two HUWE1 knockout (KO) clones of HAP1 cells. Proteins with altered abundance in response to HUWE1 knockdown are indicated in red. Proteins selected for further analysis are indicated in green. D) Immunoblots of 293T cells stably expressing FLAG-HA tagged C16orf72/HAPSTR1 were treated with bafilomycin A1 (BafA1), TAK-243, cycloheximide (CHX) or MG132 for eight hours. Whole cell extracts were separated by SDS-PAGE and immunoblotted (IB) with the indicated antibodies. E) Immunoblots of 293T cells stably expressing FLAG-HA tagged C16orf72/HAPSTR1 and transfected with a control (Ctrl) siRNA or one of three C16orf72 or HUWE1 targeting siRNAs for 72 hours. Whole cell extracts were separated by SDS-PAGE and immunoblotted (IB) with the indicated antibodies (s, short exposure; l, long exposure). F) Immunoblots of 293T cells stably expressing FLAG-HA tagged SCNM1 (top) or transiently transfected with FLAG tagged MAFB (bottom) and transfected with a control (Ctrl) siRNA or one of three HUWE1 targeting siRNAs for 72 hours. Whole cell extracts were separated by SDS-PAGE and immunoblotted (IB) with the indicated antibodies. See also Figure S1 and Tables S1-S3.

### C16orf72/HAPSTR1 is a HUWE1 substrate

One protein that accumulated substantially upon both HUWE1 knockdown and knockout was C16orf72. This protein was notable due to its known genetic and physical association with HUWE1 (Hein et al., 2015; Vazquez and Boehm, 2020). To validate C16orf72, as well as other identified proteins, as HUWE1 substrates, we generated stable FLAG-HA-tagged expression cell lines. The protein levels of stably expressed C16orf72 substantially increased upon treatment with proteasome (MG132) or ubiquitin-activating enzyme inhibitors (TAK-243), but not the autophagy inhibitor BafA1 indicating that C16orf72 is targeted for degradation by the ubiquitin-proteasome system (Figure 1D). Validating our proteomic data, HUWE1 siRNA-mediated knockdown elevated C16orf72 protein levels (Figure 1E). Similarly, HUWE1 knockdown elevated the levels of exogenous SCNM1 and MAFB, two proteins identified as putative HUWE1 substrates from our proteomic analyses (Figure 1F). HUWE1 also ubiquitylates C16orf72 *in vitro*, further substantiating C16orf72 as a HUWE1 substrate (Figure S1B). A recent study also identified C16orf72 as a HUWE1 substrate and interacting protein that broadly alters transcription of many stress-responsive proteins, renaming C16orf72 as HUWE1-Associated Protein modifying STress Responses, or HAPSTR1, which we adopt here (Amici et al., 2022).

### HUWE1 cooperates with other ligases to mediate substrate degradation

The breadth of putative HUWE1 substrates, combined with the poor overlap of identified HUWE1 substrates across cell lines, tissues, or conditions suggests that HUWE1 activity may not be restricted to dedicated substrates, but rather extends to substrates shared with other ubiquitin ligases. To begin to discriminate between dedicated versus shared HUWE1 substrates, we employed a knockdown-rescue approach. For this approach, we knocked down endogenous HUWE1 while also transiently expressing exogenous HUWE1 engineered to be resistant to the HUWE1-targeting siRNAs #2 and #3. Endogenous HUWE1 depletion stabilized both exogenous HAPSTR1 and endogenous DDIT4, a well-characterized HUWE1 substrate. Expression of exogenous, siRNA resistant HUWE1 restored HAPSTR1 and DDIT4 degradation. However, expression of a catalytically inactive version of HUWE1 containing a point mutant that replaces the catalytic cysteine residue within the HECT domain with a serine residue (C4341S, hereafter CS) failed to rescue HAPSTR1 and DDIT4 degradation (Figure 2A). We further observed that HUWE1 CS overexpression in the presence of endogenous HUWE1 stabilized both HAPSTR1 and DDIT4 (Figure 2A), suggesting that inactive HUWE1 acts in a dominant negative fashion to inhibit either wild type HUWE1 or other cooperating ubiquitin ligases that also target HAPSTR1 or DDIT4. Interestingly, expression of HUWE1 CS does not appear to act in a dominant negative manner towards HAPSTR1 when endogenous HUWE1 is depleted (Figure 2A). In contrast, HUWE1 CS expression resulted in stabilization of DDIT4 even in the absence of endogenous HUWE1 (Figure 2A). Taken together, these results suggest that HAPSTR1 is a dedicated HUWE1 substrate, and that some fraction of characterized HUWE1 substrates, including DDIT4, are shared with other ligases.

**Figure 2.**
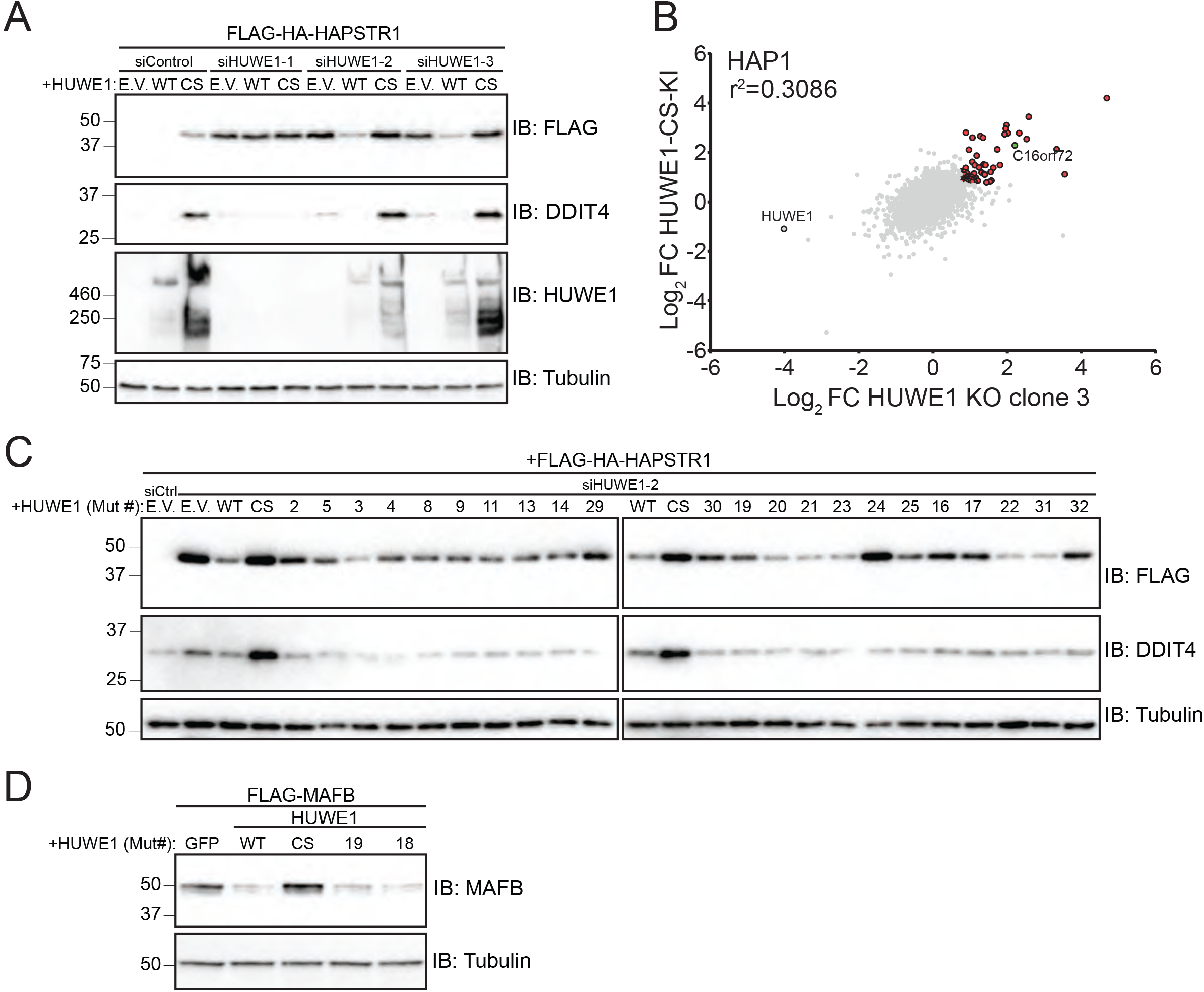
HAPSTR1 is a HUWE1 substrate. A) Immunoblots of 293T cells stably expressing FLAG-HA tagged HAPSTR1 and co-transfected with a control or HUWE1 targeting siRNA and either an empty vector (E.V.) control, wild type HUWE1 (WT), or HUWE1 C4341S (CS) for 48 hours. The HUWE1 transgenes are resistant to siHUWE1-2 and siHUWE1-3. Whole cell extracts were separated by SDS-PAGE and immunoblotted (IB) with the indicated antibodies. B) Scatter plot comparing the log_2_ fold change (FC) for all proteins quantified by tandem mass tag proteomics in a HUWE1 knockout HAP1 clone and a HAP1 clone with the endogenous HUWE1 locus edited with a cysteine to serine mutation at residue 4341 (CS). Proteins whose abundance is significantly elevated in both cell lines are shown in red. Proteins selected for further analysis are indicated in green. C) Immunoblots of 293T cells stably expressing FLAG-HA tagged HAPSTR1 and co-transfected with a control (siCtrl) or HUWE1-targeting siRNA and either an empty vector (E.V.) control, or the indicated variant of siRNA resistant HUWE1 (WT, wild type; CS, C4341S). See Table S4 for mutation details for the indicated HUWE1 variants. Whole cell extracts were separated by SDS-PAGE and immunoblotted (IB) with the indicated antibodies. D) Immunoblots of 293T cells transiently transfected with FLAG tagged MAFB and either a GFP control or the indicated HUWE1 variant (WT, wild type; CS, C4341S). See Table S4 for mutation details for the indicated HUWE1 variants. Whole cell extracts were separated by SDS-PAGE and immunoblotted (IB) with the indicated antibodies. See also Figure S2 and Table S4.

Our observation that expression of inactive HUWE1 stabilized substrates in a manner distinct from HUWE1 loss of function suggested a possible strategy to identify a larger population of HUWE1 substrates. We used genome engineering to generate a HUWE1 knock-in (KI) HAP1 cell line that replaces the catalytic cysteine with serine at the genomic locus. We analyzed proteome changes in this HAP1 HUWE1 knock-in cell line compared to parental HAP1 cells using our quantitative proteomic pipeline. Of the more than 7300 proteins quantified, 710 displayed altered abundance with 427 increased abundance. As expected, HAPSTR1 levels were elevated in the HUWE1 KI cells (Figure 2B; Table S3). The larger number of putative substrates identified in the HUWE1 KI cells compared to HUWE1 KO cells suggests that our strategy to identify a larger pool of HUWE1 substrates was effective, and that many of the substrates identified in HUWE1 KI cells, but not in the HUWE1 knockout cells, may be shared substrates with other ligases. Of the identified proteins with elevated protein levels in both HUWE1 KO clones, 66% also had increased protein abundance in HUWE1 knock-in cells. Despite this increase in substrate identification, only 17% (19/107) of the initial HUWE1 substrates identified using shRNA in 293T cells also displayed elevated protein levels in HAP1 HUWE1 knock-in cells. This observation again suggests that HUWE1 substrates may be largely tissue or cell type-specific. Consistent with this observation, comparing all proteomics experiments in HUWE1 loss of function cells across three cell types reveals that more than 80% of all putative HUWE1 substrates identified in this study were identified in a single experiment with only a single common substrate shared among the four experiments (Figures S2A-C). This observation may explain why previously characterized HUWE1 substrates are largely absent in our experiments, instead identifying a new pool of putative HUWE1 substrates. Combined, our data suggest that HUWE1 targets a broad, and highly cell type-specific substrate pool, which may be a desired feature for a plastic ubiquitin ligase tasked to broadly regulate disparate pathways and substrates.

### HUWE1 structural features differentially enable substrate degradation

We used our knockdown rescue approach with a collection of HUWE1 mutants to start to understand which HUWE1 sequence and structural features are utilized for substrate targeting. We generated over 30 siRNA resistant HUWE1 variants containing either mutations in key domains or structural features, or disease-associated mutations, and we tested the ability of each variant to destabilize HAPSTR1 and DDIT4. (Figure S3A; Table S4). HUWE1 contains two separate ubiquitin-binding modules, one consisting of a UIM and a UBA domain, and one built from three repetitive UBM motifs (Hunkeler *et al.*, 2021). HUWE1 variants lacking individual ubiquitin-binding entities were partially compromised in their ability to target HAPSTR1 for degradation compared to wild type re-expression. HAPSTR1 levels were further stabilized by expression of combinatorial mutants eliminating multiple ubiquitin-binding domains suggesting that the ability of HUWE1 to bind ubiquitin contributes to HAPSTR1 degradation (Figures 2C and S3B). A panel of five HUWE1 mutants observed in patients with intellectual disability disorders largely resembled wild type HUWE1 in its ability to target HAPSTR1 with only the larger Δ168-189 patient-derived variant compromising HUWE1 function. Two HUWE1 structural mutants (#24 and #32) were also unable to destabilize HAPSTR1, with the deletion of a large unstructured domain impairing HAPSTR1 degradation to a level only observed with the catalytically inactive (CS) variant. Interestingly, neither the ubiquitin-binding domains nor the large unstructured domain within HUWE1 were required for HUWE1 to target DDIT4 (Figure 2C). Similarly, a HUWE1 variant lacking all ubiquitin-binding domains retained its ability to target exogenous MAFB (Figures 2D and S3B). These results suggest that diverse structural features within HUWE1 operate in a substrate-specific manner.

### A subset of HUWE1 substrates requires HAPSTR1

Examination of DepMap data, which examines cell proliferation upon CRISPR-based knockout of individual genes across a panel of >500 cell lines (Behan et al., 2019; Dempster et al., 2019; Vazquez and Boehm, 2020), revealed a strong positive genetic correlation between HUWE1 and HAPSTR1 as noted by others (Amici et al., 2022) (Figure 3A). This relationship was not expected, given our clear demonstration that HAPSTR1 is a HUWE1 substrate, as ligase-substrate relationships would be predicted to have a negative correlation. This observation suggests that HAPSTR1 is both a substrate and a positive regulator of HUWE1 function. To gain insights into how HAPSTR1 regulates HUWE1 function, we measured protein abundance changes upon knockdown of HUWE1 or HAPSTR1 using our unbiased quantitative proteomic pipeline. We utilized CAL27 cells for this approach as DepMap studies identified CAL27 cells to be dependent on both HUWE1 and HAPSTR1 for proliferation. We identified 115 and 106 proteins whose protein abundance increased in response to HUWE1 or HAPSTR1 knockdown, respectively (Figure 3B; Table S5). Consistent with previous results, putative HUWE1 substrates identified in CAL27 cells were largely unique and not identified as HUWE1 substrates in our studies in other cell lines, or in previous studies (Figures S2A-C). Interestingly, protein abundance changes comparing HUWE1 and HAPSTR1 knockdown were significantly correlated with 40% of putative HUWE1 substrates also accumulating upon HAPSTR1 knockdown (Figure 3B; Table S5). To validate these results, we generated CAL27 cells with stable expression of a subset of identified substrates. Despite not identifying HAPSTR1 in our CAL27 proteomic study, we confirmed that HAPSTR1 is a robust HUWE1 substrate in CAL27 cells (Figure 3C). We also validated SCNM1, MAFB, NFIB, and ZCCHC17 as HUWE1 substrates that are also stabilized upon HAPSTR1 knockdown (Figure 3C). We noted variation in the dependence of some substrates on HUWE1 versus HAPSTR1. For example, NFIB was stabilized to a greater extent upon HAPSTR1 knockdown compared to HUWE1 knockdown and MAFB was more dependent on HUWE1 than HAPSTR1. These results suggest that substrates utilize HUWE1 or HAPSTR1 to different extents, arguing that both factors may operate independently from each other in some contexts. Regardless, our data implicate HAPSTR1 as a putative regulator of HUWE1 function and identify HUWE1 substrates that also require HAPSTR1 for degradation.

**Figure 3.**
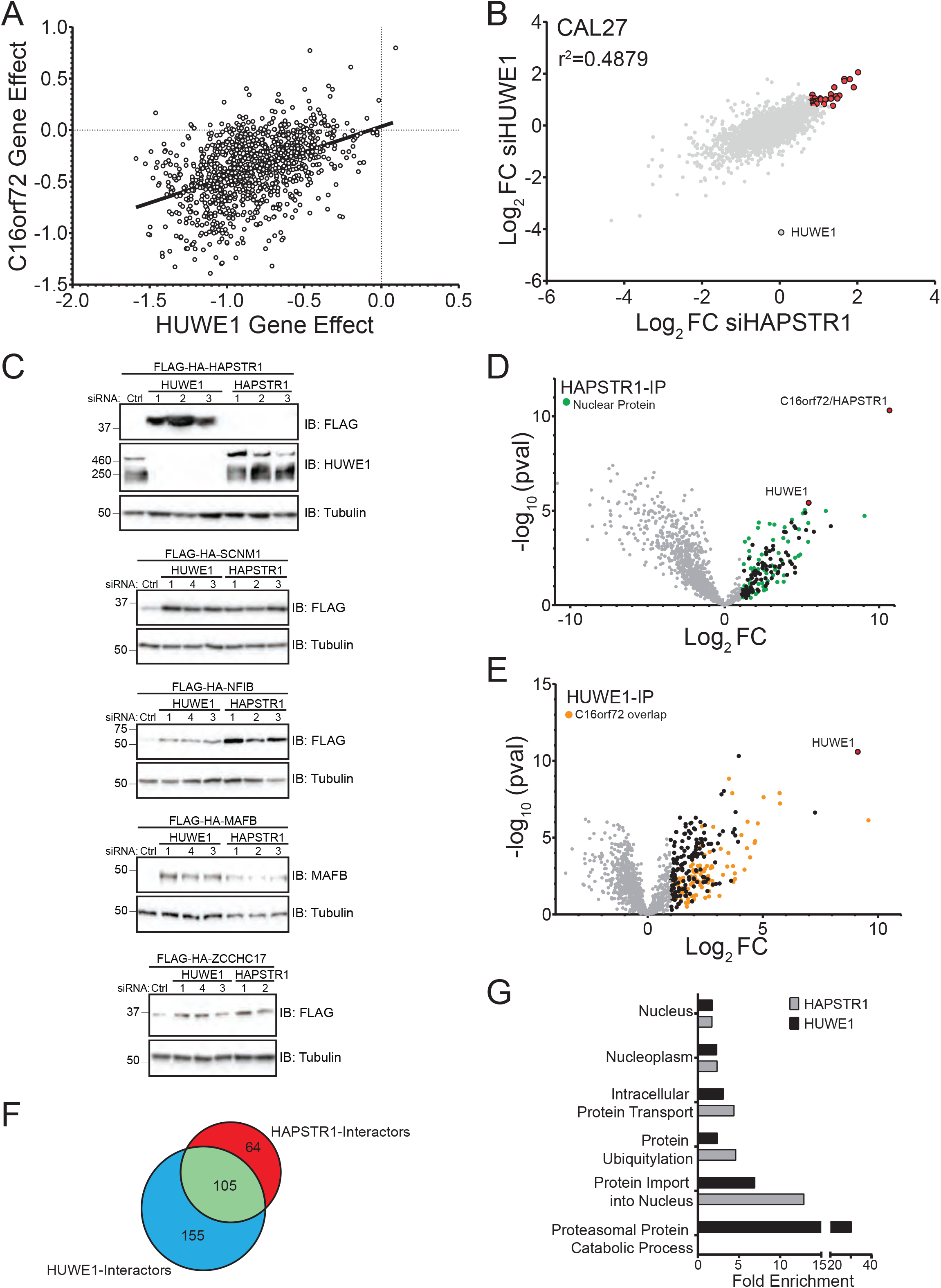
HAPSTR1 assists HUWE1 in targeting nuclear substrates. A) Scatter plot of DepMap 22Q2 CRISPR gene effect scores for HUWE1 and C16orf72/HAPSTR1. The solid line depicts the best-fit line using linear regression. B) Scatter plot comparing the log_2_ fold change (FC) for all quantified proteins in CAL27 cells 72 hours after transfection of a HUWE1 or HAPSTR1 targeting siRNA relative to a control siRNA. Proteins with increased abundance in both HUWE1 and HAPSTR1 knockdown cells are indicated in red. C) Immunoblots of CAL27 cells stably expressing the indicated FLAG-HA tagged transgene and transfected with siRNA for 72 hours. Whole cell extracts were separated by SDS-PAGE and immunoblotted (IB) with the indicated antibodies. D) FLAG-tagged HAPSTR1 was transiently expressed and HAPSTR1 associated proteins were identified by immunoprecipitation followed by mass spectrometry (IP-MS). The volcano plot depicts the log_2_ fold change (FC) and -log_10_ P-value for proteins identified in HAPSTR1 immune complexes compared to control IPs. Proteins that are significantly enriched with HAPSTR1 are shown as black dots. Significantly enriched proteins with reported nuclear localization are depicted as green dots. E) FLAG-tagged HUWE1 was transiently expressed and associated proteins were identified by IP-MS. The volcano plot depicts the log_2_ fold change (FC) and -log_10_ P-value for proteins identified in HUWE1 immune complexes compared to control IPs. Proteins that are significantly enriched with HUWE1 are shown as black dots. HUWE1 interacting proteins that are also enriched with HAPSTR1 are depicted as orange dots. F) Overlap of identified HUWE1 and HAPSTR1 interacting proteins. G) Fold enrichment of indicated gene ontology terms associated with HUWE1 or HAPSTR1 interacting proteins. See also Figure S3 and Tables S5 and S6.

### HAPSTR1 interacts with HUWE1 and nuclear proteins

Given the positive genetic relationship between HUWE1 and HAPSTR1, and our demonstration that some HUWE1 substrates require HAPSTR1 for targeting, we postulated that HAPSTR1 may act as a substrate adaptor for HUWE1. We therefore mapped the physical interactions for each protein using affinity-enrichment followed by mass spectrometry. We immunoprecipitated FLAG tagged HAPSTR1 or HUWE1 with anti-FLAG antibodies and compared the identified proteins to those identified from FLAG-GFP isolations. Consistent with previous studies (Amici et al., 2022; Benslimane et al., 2021; Hein et al., 2015), HAPSTR1 and HUWE1 interact with each other (Figures 3D and 3E). Further, more than 60% of the identified HUWE1 interacting proteins were also identified as HAPSTR1 interacting proteins, with 40% of HAPSTR1 interactors overlapping with HUWE1 interactors (Figure 3F; Table S6). This substantial degree of shared interacting proteins is consistent with the hypothesis that HAPSTR1 associates with HUWE1 to target common substrates or that HAPSTR1 and HUWE1 engage a common set of regulators. Interestingly, bioinformatic analyses of HAPSTR1- and HUWE1-interacting proteins revealed an enrichment for nuclear localized proteins and for proteins known to regulate nuclear cytoplasmic transport (Figure 3G). HUWE1 interacting proteins additionally displayed enrichment for proteins involved in proteasomal degradation, including many proteasomal proteins, as previously described (Tai et al., 2010). However, these proteasomal proteins were notably absent from the list of HAPSTR1 interactors. This observation, and the finding that HUWE1 protein levels far exceed HAPSTR1 in nearly all cell and tissue types (Nusinow et al., 2020; Thul et al., 2017; Uhlen et al., 2015), suggests that HUWE1 likely functions in complexes with and without HAPSTR1. In total, while these data are consistent with the idea that HAPSTR1 functions as a HUWE1 substrate adaptor, only a defined subset of HUWE1 likely cooperates with HAPSTR1.

### HAPSTR1 localizes HUWE1 to the nucleus

Combining our interaction data indicating that HAPSTR1 and HUWE1 both engage nuclear proteins with our identification and validation of nuclear proteins that are stabilized by either HAPSTR1 or HUWE1 loss of function, we speculated that HAPSTR1 may regulate nuclear HUWE1 functions. To test this hypothesis, we first knocked down HUWE1 or HAPSTR1 and biochemically isolated nuclei. Subsequent immunoblotting for endogenous HUWE1 revealed that HAPSTR1 knockdown substantially reduced nuclear HUWE1 levels, resulting in cytoplasmic enrichment (Figures 4A, S4A). Consistent with our previous proteomics results, HAPSTR1 knockdown does not change total HUWE1 protein levels (Figure 3B and Table S5). These data suggest that HAPSTR1 plays a role in localizing HUWE1 to the nucleus.

**Figure 4.**
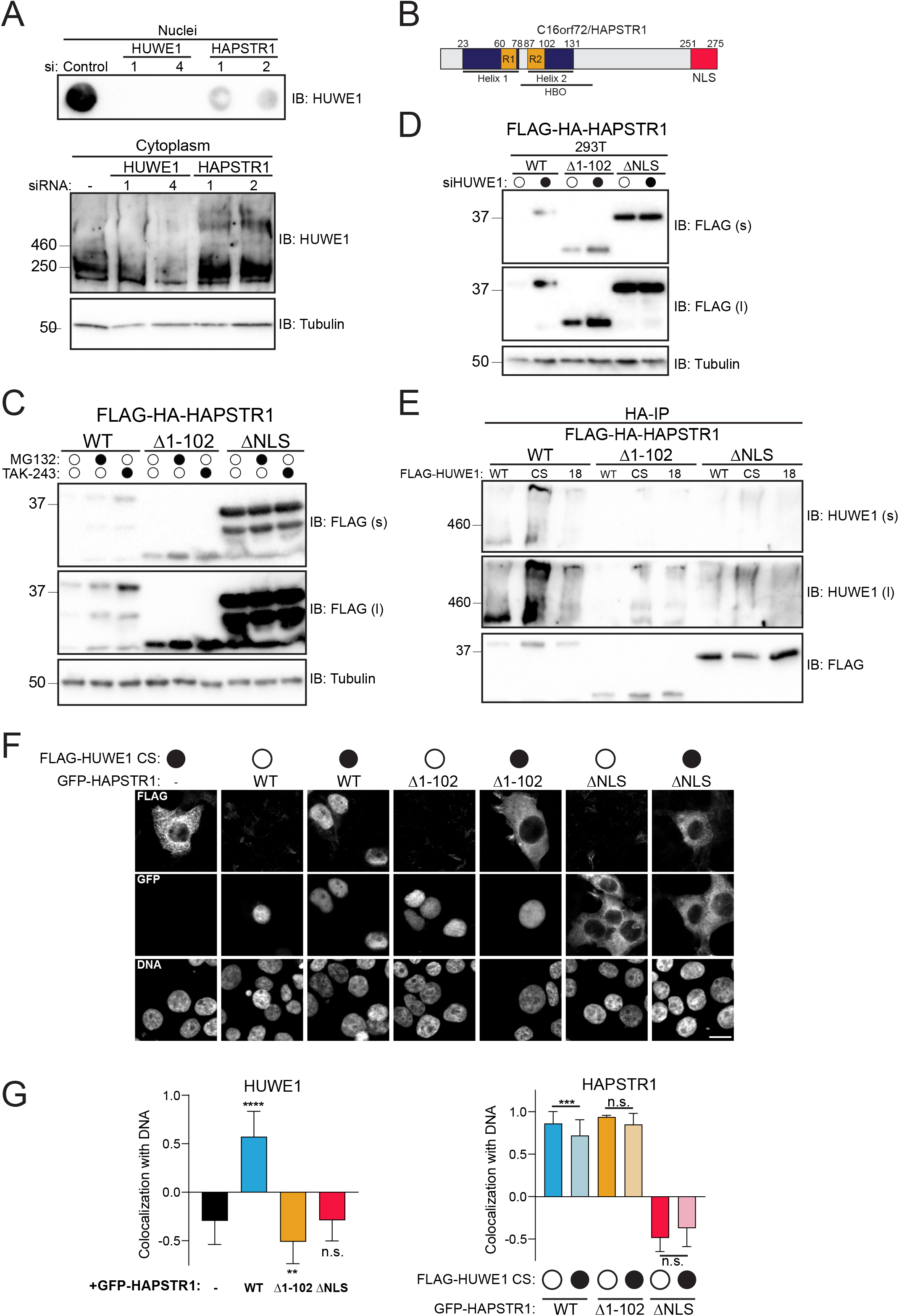
HUWE1 nuclear localization requires HAPSTR1. A) Immunoblots of NCI-H2052 cells transfected with a control siRNA or one of two siRNA sequences targeting either HUWE1 or HAPSTR1 for 72 hours. (Top) The nuclei were isolated and analyzed by dot blot. (Bottom) The cytoplasmic fractions were separated by SDS-PAGE. Immunoblots (IB) were performed with the indicated antibodies. B) Schematic of HAPSTR1 protein sequence features. Predicted alpha helices are shown in purple, conserved repeated regions in orange, and nuclear localization sequence in red. C) Immunoblots of 293T cells stably expressing FLAG-HA tagged wildtype (WT) or the indicated variant of HAPSTR1 treated with MG132 or TAK-243 for eight hours. Whole cell extracts were separated by SDS-PAGE and immunoblotted (IB) with the indicated antibodies. D) Immunoblots of 293T cells stably expressing siRNA resistant FLAG-HA tagged wildtype (WT) or the indicated variant of HAPSTR1 transfected with a control or HUWE1-targeting siRNA. Whole cell extracts were separated by SDS-PAGE and immunoblotted (IB) with the indicated antibodies (s, short exposure; l, long exposure). E) Immunoblots of HA immunoprecipitations from 293T cells stably expressing FLAG-HA tagged wildtype (WT) or the indicated variant of HAPSTR1 and transfected with the indicated FLAG tagged HUWE1 variant (WT, wild type; CS, C4341S; 18, deletion of all UBA, UIM, and UBM domains). HA-IP eluates were separated by SDS-PAGE and immunoblotted (IB) with the indicated antibodies (s, short exposure; l, long exposure). F) Immunofluorescence images of 293T cells transfected with FLAG-tagged HUWE1 C4341S (CS) alone or in combination with the indicated GFP tagged HAPSTR1. Scale bar, 20 μm. G) Quantification of the colocalization of DNA and HUWE1 (left) or HAPSTR1 (right). Data are represented as the mean + SD for all cells quantified. Analysis included at least two biological replicates for all conditions. Statistical significance was determined by Kolmogorov-Smirnov t-test. **** p ≤ 0.0001; ** p ≤ 0.01; n.s. p > 0.05. See also Figure S4.

We next sought to define HAPSTR1 mutations that perturb either the HUWE1 interaction, or the nuclear localization of HAPSTR1. Sequence comparison of human HAPSTR1 with its putative ortholog in saccharomyces cerevisiae (YJR056C) identified a repeated conserved N-terminal sequence this is contained within two predicted helical domains (Figure 4B). Recent studies mapped a HUWE1 interaction domain (HBO) within this region (Amici *et al.*, 2022), and a putative nuclear localization sequence (NLS) was identified in the C-terminus of HAPSTR1. We therefore generated two HAPSTR1 deletion mutants, one without the first 102 amino acids that includes the N-terminal conserved repeated regions (R1 and R2; Δ1-102) and one with the C-terminal NLS deleted (ΔNLS). Stable 293T cell lines expressing wild type and the two HAPSTR1 mutants demonstrated elevated HAPSTR1 protein levels for the Δ1-102 variant compared to wild type and a further striking increase in protein levels upon NLS deletion (Figures 4C and S4B). Both wild type and Δ1-102 HAPSTR1 protein levels increased upon proteasome or ubiquitin activating enzyme inhibition as well as upon HUWE1 knockdown, although the extent of stabilization for Δ1-102 was less than wild type HAPSTR1 (Figures 4C, 4D, and S4B). In contrast, the ΔNLS HAPSTR1 variant was not stabilized by proteasome inhibition or HUWE1 knockdown (Figures 4C, 4D, and S4B). While similar results were observed in CAL27 cells with stable expression of wild type or mutant HAPSTR1, Δ1-102 HAPSTR1 was expressed at levels lower than observed in 293T cells (Figures S4C-E). Wild type HAPSTR1 interacted with HUWE1 in co-IP experiments and this interaction was substantially compromised, but not eliminated, upon deletion of either the N-terminal 102 amino acids or the C-terminal NLS (Fig. 4E). This result suggests that, while the N-terminus contributes to HUWE1 binding, sequences beyond the repeated regions, like the larger HBO domain, as well as the C-terminal NLS, contribute critical HUWE1 binding surfaces.

To further test if HAPSTR1 localizes HUWE1 to the nucleus, we examined exogenous HUWE1 cellular localization by microscopy with and without HAPSTR1 co-expression. HUWE1 was largely localized to the cytoplasm while HAPSTR1 was nuclear when expressed in isolation (Figures 4F and 4G). Upon co-expression, wild type HAPSTR1 strikingly relocalized co-expressed HUWE1 to the nucleus (Figures 4F and 4G). HUWE1 co-expression with the Δ1-102 variant of HAPSTR1 did not result in HUWE1 nuclear localization, despite Δ1-102 HAPSTR1 maintaining its nuclear localization. As expected, ΔNLS HAPSTR1 was relocalized to the cytoplasm and failed to enrich HUWE1 within the nucleus (Figures 4F and 4G). In total, our results demonstrate that HAPSTR1 is required for nuclear localization of HUWE1.

### HAPSTR1 potentiates nuclear HUWE1 activity

Because nuclear HUWE1 levels are regulated by HAPSTR1, we examined if the ability of HUWE1 to target nuclear substrates was similarly regulated by HAPSTR1. We used a knockdown rescue approach in cell lines with stable expression of siRNA-resistant, FLAG-HA tagged wild type, Δ1-102, or ΔNLS HAPSTR1. We then expressed two substrates, MAFB and SCNM1, that we previously demonstrated to be stabilized by either HUWE1 or HAPSTR1 knockdown (Figure 3C) in these HAPSTR1 expressing cell lines with or without knockdown of endogenous HAPSTR1. HAPSTR1 knockdown elevated MAFB and SCNM1 protein levels in parental lines, as expected, and re-expression of wild type HAPSTR1 largely eliminated the increase in MAFB or SCNM1 levels upon endogenous HAPSTR1 knockdown (Figures 5A and 5B). The Δ1-102 and the ΔNLS HAPSTR1 variants did not rescue MAFB nor SCNM1 degradation, with the ΔNLS version acting in a dominant negative fashion with regards to MAFB degradation (Figures 5A and 5B). These results suggest that loss of HAPSTR1 restricts HUWE1 localization to the cytoplasm, thereby impacting HUWE1’s ability to target nuclear substrates.

**Figure 5.**
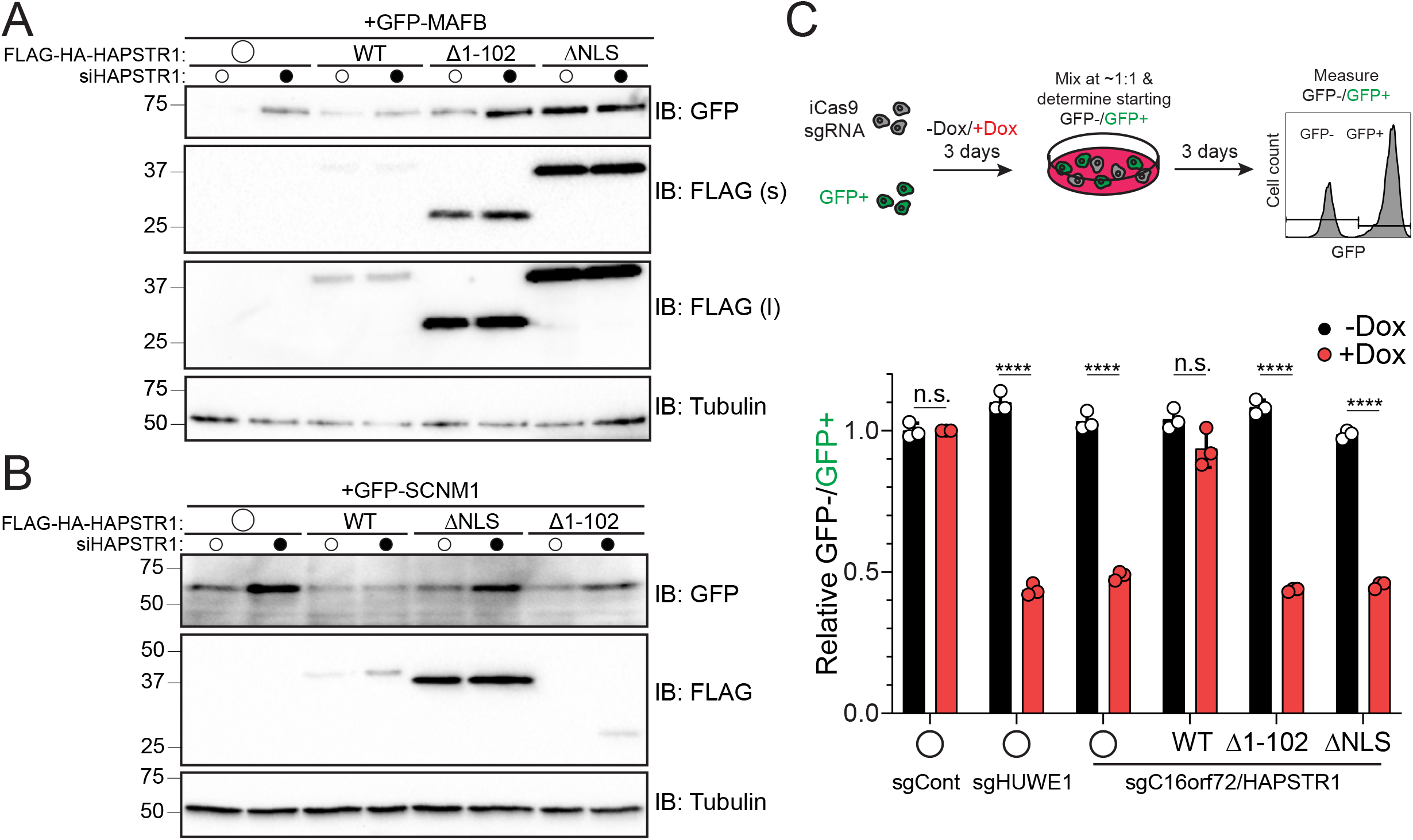
Loss of nuclear HUWE1 stabilizes nuclear targets and reduces cellular proliferation. A) Immunoblots of 293T cells stably expressing the indicated siRNA resistant FLAG-HA-tagged HAPSTR1 variant and transfected with GFP-tagged MAFB and a control or HAPSTR1 targeting siRNA for 72 hours. Whole cell extracts were separated by SDS-PAGE and immunoblotted (IB) with the indicated antibodies (s, short exposure; l, long exposure). B) Immunoblots of CAL27 cells stably expressing GFP-SCNM1 and the indicated variant of siRNA resistant FLAG-HA tagged HAPSTR1 and transfected with a control or HAPSTR1 targeting siRNA for 72 hours. Whole cell extracts were separated by SDS-PAGE and immunoblotted (IB) with the indicated antibodies. C) Top, schematic of the internally controlled CRISPR-Cas9-based competitive growth assay. Bottom, CAL27 cells expressing inducible Cas9 and control, HUWE1, or HAPSTR1 targeting sgRNA alone or with stable expression of the indicated sgRNA resistant HAPSTR1 variant were untreated or dox treated for 72hrs prior to mixing with GFP expressing parental CAL27 cells. The relative Non-GFP:GFP ratio of the population was measured 72hrs after co-culture. Data are represented as the mean ± SD of three replicates. Statistical significance was determined by unpaired t-test. **** p ≤ 0.0001; n.s. p > 0.05. See also Figure S5.

We next wondered if nuclear HUWE1 activity is required to support overall cell proliferation. We generated CAL27 cells with doxycycline-inducible Cas9 expression and stable expression of an sgRNA targeting HUWE1 or HAPSTR1. We confirmed that Cas9 expression in cells expressing a HUWE1-targeting sgRNA resulted in decreased HUWE1 protein levels (Figures S5A and S5B). To determine the effect of HUWE1 or HAPSTR1 knockout on cellular proliferation, we used a competitive growth assay in which control GFP expressing cells were mixed 1:1 with iCas9 expressing cells (Figure 5C). The nonGFP:GFP cell ratio was then measured over time. Expression of an sgRNA targeting HUWE1 or HAPSTR1, but not the control sgRNA, resulted in a doxycycline-dependent decrease in cell proliferation (Figure 5C). This result validates the DepMap data showing a loss of CAL27 cell proliferation upon HUWE1 or HAPSTR1 knockout. We then re-expressed CherryFP-tagged WT, Δ1-102, or ΔNLS HAPSTR1 in the inducible Cas9 CAL27 cells (Figure S5A). The sgRNA targeting HAPSTR1 used here targets the 5’UTR and start codon of the endogenous HAPSTR1 locus, a sequence which is not present in our transgene. Expression of wild type, but not the Δ1-102 or ΔNLS HAPSTR1 variants rescued the dox-inducible repression of cell proliferation upon targeting of the endogenous HAPSTR1 locus (Figure 5C). These results argue that loss of nuclear HUWE1 represses cell proliferation in CAL27 cells. These results also indicate that the growth suppression observed upon HAPSTR1 loss of function likely stems from a loss of nuclear HUWE1 levels.

### Nuclear HUWE1 broadly regulates a stress-dependent transcriptional program

To more broadly examine the role of nuclear HUWE1, we performed RNA-sequencing in CAL27 cells after siRNA-mediated knockdown of HUWE1 or HAPSTR1. Loss of either HUWE1 or HAPSTR1 resulted in broad transcriptional changes with 845 and 677 significantly differentially expressed genes observed upon HUWE1 or HAPSTR1 knockdown, respectively (Figure 6A-C, and S6A; Table S7). We noted a correlation between the altered transcripts upon HUWE1 or HAPSTR1 knockdown, with 405 shared differentially expressed genes, suggesting that loss of HAPSTR1 or HUWE1 impacts similar transcriptional pathways (Figures 6A-C, and S6B). The majority of observed protein level changes in our CAL27 proteomics dataset were not observed on the transcript level, although some protein level changes for noted HUWE1 substrates were likely the combined result of altered protein turnover and increased mRNA abundance (Figures S6C and S6D). Pathway enrichment analysis of differentially expressed genes revealed repression of many genes involved in innate immunity and inflammatory signaling upon loss of HUWE1 or HAPSTR1 (Figures 6D and 6E). Transcriptional regulatory pathways were enriched in transcripts that both increased and decreased upon HUWE1 knockdown, suggesting a high degree of variability in the mechanism by which HUWE1 impacts transcription (Figures 6D and 6E). These results are broadly consistent with a recent study demonstrating that similar stress-responsive transcriptional pathways are impacted by either loss of HUWE1 or HAPSTR1 (Amici et al., 2022).

**Figure 6.**
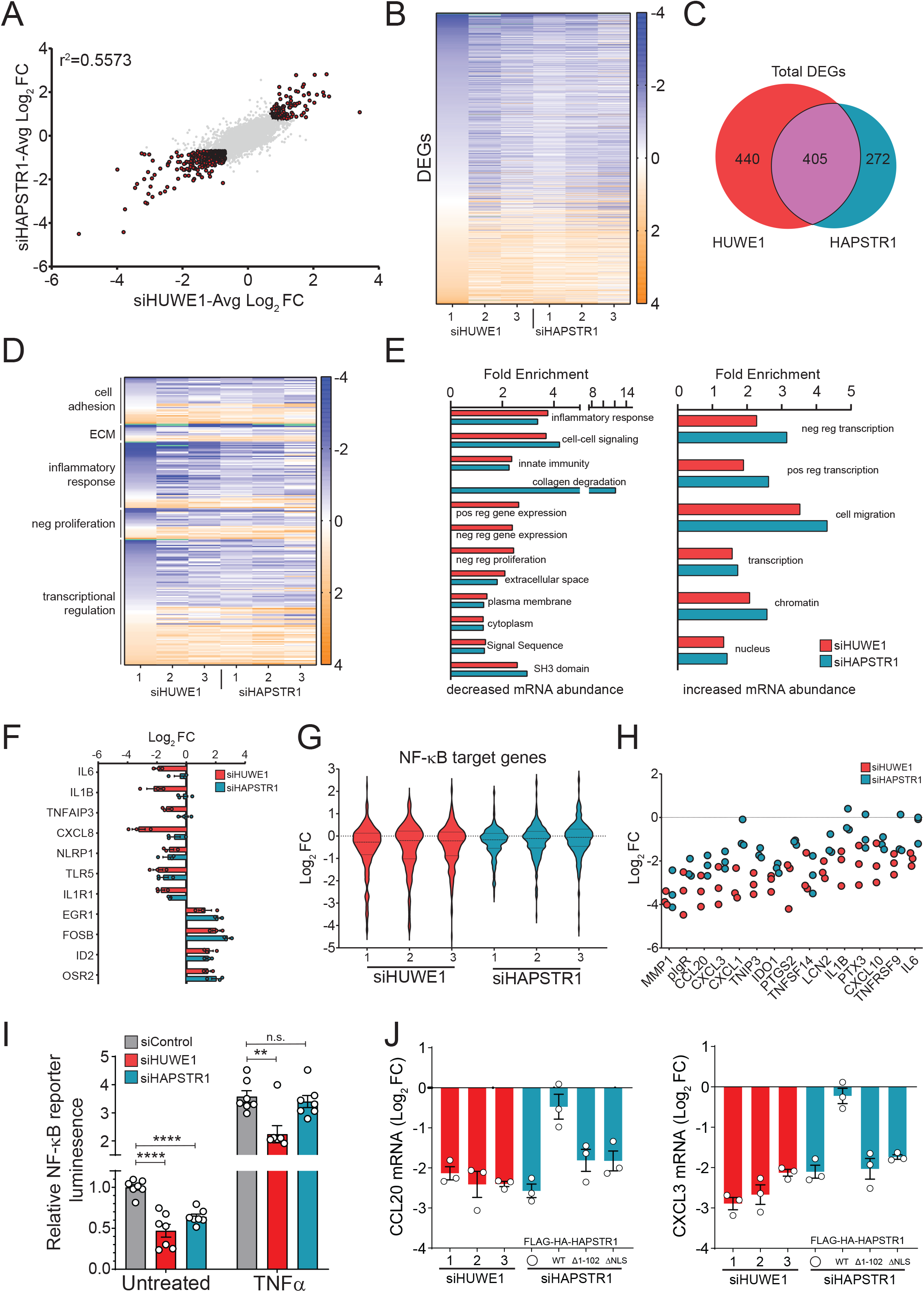
HAPSTR1 and HUWE1 regulate overlapping transcriptional programs. A) Scatter plot comparing the log_2_ fold change (FC) for all mRNA quantified by RNA-seq in CAL27 cells 72 hours after transfection of a HUWE1 or HAPSTR1 targeting siRNA relative to a control siRNA. The data represent the average across three different siRNA sequences targeting HUWE1 and HAPSTR1. Differentially expressed genes in both HUWE1 and HAPSTR1 knockdown cells are indicated in red. B) Heat map of the differentially expressed genes after HUWE1 or HAPSTR1 knockdown in CAL27 cells. Color map indicates log_2_ fold change. C) Venn diagram of the differentially expressed genes after HUWE1 or HAPSTR1 knockdown in CAL27 cells. D) Heat map of a subset of differentially expressed genes after HUWE1 or HAPSTR1 knockdown in CAL27 cells clustered by selected enriched gene ontology terms. E) Selected enriched gene ontology biological processes, cellular components, or pfam domains of mRNAs with decreased abundance (left) or increased abundance (right) in CAL27 cells after HUWE1 or HAPSTR1 knockdown. F) The log_2_ fold change (FC) of selected inflammatory or transcriptional regulation genes in CAL27 cells after HUWE1 or HAPSTR1 knockdown. N=3. Error bars = SD G) Violin plot depicting the log_2_ fold changes of all identified NF-κB target genes in CAL27 cells after HUWE1 or HAPSTR1 knockdown by three different siRNA target sequences each. H) Log_2_ fold changes of select NF-κB target genes in CAL27 cells after HUWE1 or HAPSTR1 knockdown. Each dot represents the mean of three replicates per siRNA used. I) 293T cells were transfected with siRNAs targeting the indicated genes followed by transfection with a NF-κB transcriptional reporter. Cells were either untreated or treated with 100 ng/mL TNFα for 6 h before luciferase activity analysis. Data are represented as the mean ± SD of four replicates. Statistical significance was determined by unpaired t-test. **** p ≤ 0.0001; ** p ≤ 0.01; n.s. p > 0.05. J) CAL27 cells were transfected with control or siRNAs targeting HUWE1 or HAPSTR1 in parental cells or cells with stable expression of the indicated siRNA-resistant HAPSTR1 variant. The abundance of *CCL20* and *CXCL3* mRNA in each sample normalized to that of *GAPDH* was determined by qRT-PCR analysis. Values in transgene-expressing cell lines were normalized to siControl. Data are represented as the mean ± SEM of three biological replicates. See also Figure S6 and Table S7.

The noted enrichment of inflammatory genes that were repressed upon HUWE1 or HAPSTR1 knockdown suggested that nuclear HUWE1 may potentiate activation of innate immune signaling pathways (Figures 6E and 6F). Mining the list of altered transcripts for enrichment of transcription factor binding sites revealed NF-κB target genes to be strongly represented within the differentially expressed genes (Figure 6G) (Roider et al., 2009). Examination of all known NF-κB targets revealed a significant repression amongst a broad range of NF-κB targets upon HUWE1 knockdown, and to a lesser extent HAPSTR1 knockdown (Figure 6G and 6H). We then turned to an NF-κB reporter system to directly test if HUWE1 or HAPSTR1 knockdown repressed baseline, or TNFα stimulated NF-κB signaling (Zhou et al., 2018). Firefly luminescence from a transfected NF-κB transcriptional reporter was increased ~3.5 fold upon TNFα stimulation (Figure 6I). Knockdown of either HUWE1 or HAPSTR1 repressed baseline NF-κB transcriptional activity with HUWE1 but not HAPSTR1 impacting the TNFα stimulated response (Figure 6I). These results are consistent with previous studies demonstrating that loss of HUWE1 suppresses NF-κB activity (Ohtake et al., 2016). We then used qPCR to validate that CCL20 and CXCL3 mRNA expression was repressed in unstimulated CAL27 cells upon knockdown of either HUWE1 or HAPSTR1 (Figure 6J). Using CAL27 cell lines with stable expression of wild type and mutant HAPSTR1, we demonstrated that the observed repression of CCL20 and CXCL3 expression upon HAPSTR1 knockdown was reversed upon re-expression of wild type, but not Δ1-102 or ΔNLS, HAPSTR1 variants (Figure 6J). These results indicate that loss of nuclear HUWE1 activity is likely responsible for the observed transcriptional repression of inflammatory signaling components. Taken together, our results demonstrate that nuclear HUWE1 potentiates NF-κB activity under basal conditions and may widely regulate inflammatory signaling and responses.

### Loss of nuclear HUWE1 activates p53 signaling and limits cell proliferation

A closer examination of the DepMap data revealed a significant difference in the degree of cell proliferation repression following HUWE1 or HAPSTR1 knockout comparing cells with wild type p53 to cells containing “hotspot” p53 mutations known to inactivate p53 transcriptional activity (Figure 7A). Comparing DepMap correlations between HUWE1, HAPSTR1, and known p53 positive and negative regulators revealed a similarity between MDM2, a known negative regulator of p53 signaling, and HAPSTR1 (Figures 7A and 7B). These observations, along with previous studies, suggest that HAPSTR1 may negatively regulate p53 signaling (Benslimane et al., 2021; Chen et al., 2005; Hao et al., 2012; Kon et al., 2012; Yang et al., 2018). The noted absence of any p53 transcriptional activation upon HUWE1 or HAPSTR1 knockdown in our RNA-seq studies likely reflects that 293T and CAL27 cells lack functional p53 signaling. To directly test if HUWE1 or HAPSTR1 negatively regulate p53 signaling, we knocked down HUWE1 or HAPSTR1 in p53 functional HCT116 or RPE1 cells. Consistent with previous studies, both HUWE1 and HAPSTR1 knockdown increased p53 levels (Figures 7C, S7A-C) (Benslimane et al., 2021; Chen et al., 2005). Because HUWE1 has been reported to both positively and negatively regulate DNA damage response or repair pathways, we investigated if loss of HUWE1 or HAPSTR1 activated DNA damage response pathways which would then activate p53 signaling. Despite a clear induction of DNA damage response pathways upon etoposide or cisplatin treatment in p53 wildtype or null HCT116 cells (Figure S7A), loss of HUWE1 or HAPSTR1 did not result in a basal increase in γH2AX phosphorylation. These results suggest that loss of HUWE1 or HAPSTR1 may directly impact p53 signaling independent of the DNA damage response.

**Figure 7.**
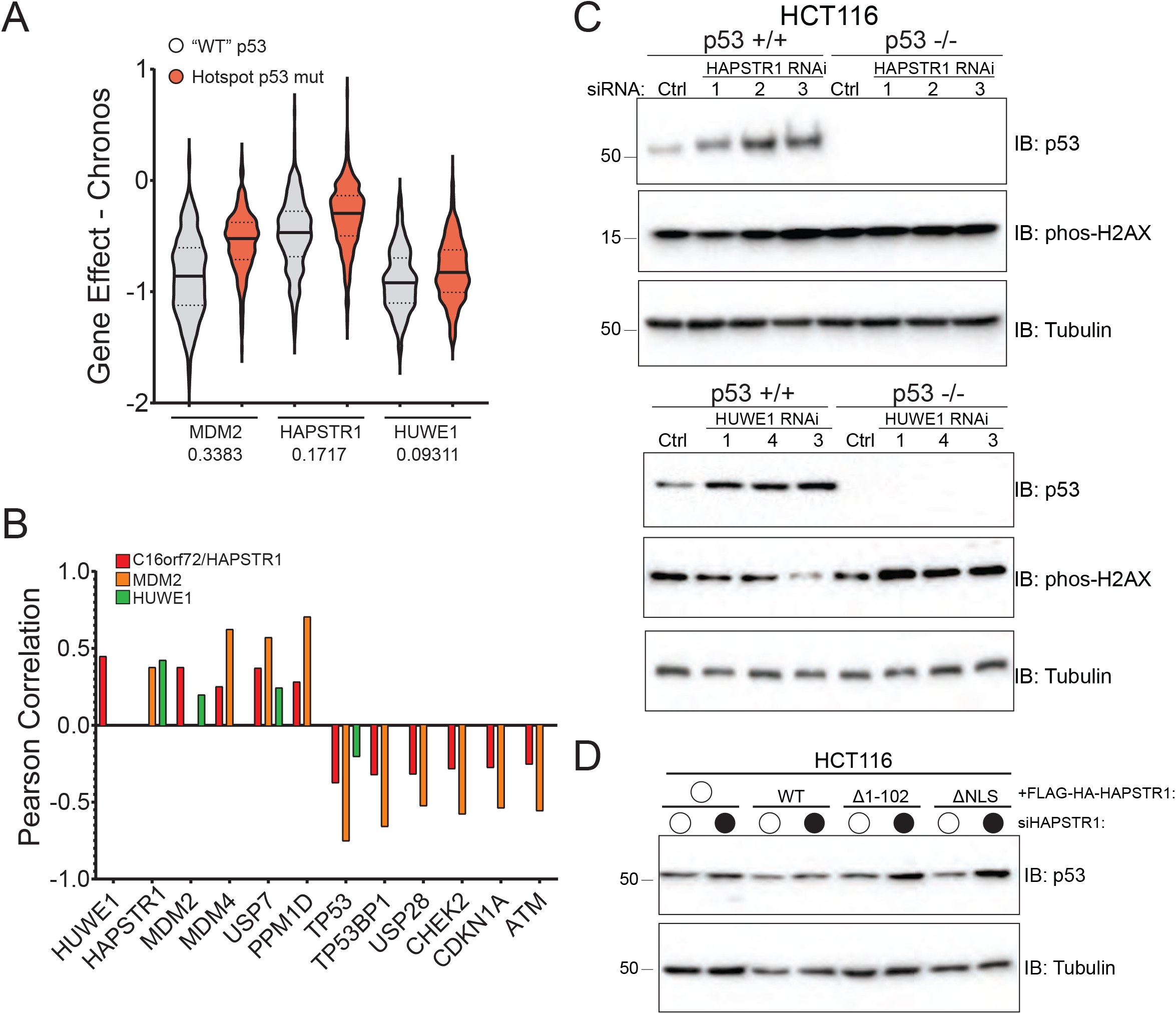
Loss of nuclear HUWE1 activates p53 signaling. A) Violin plot of DepMap 22Q2 CRISPR gene effect scores for MDM2, HAPSTR1, and HUWE1 in cell lines with wild type or hotspot mutations in p53. Numbers below gene name indicate mean difference comparing p53 wild type or hotspot mutant cell lines. B) Pearson correlation values from DepMap 22Q2 CRISPR data comparing MDM2, HAPSTR1, and HUWE1 to the indicated p53 target gene or regulator. C) Immunoblots of HCT116 cells with the indicated p53 status transfected with a control (Ctrl) siRNA or one of three siRNA sequences targeting HAPSTR1 (top) or HUWE1 (bottom). Whole cell extracts were separated by SDS-PAGE and immunoblotted (IB) with the indicated antibodies. D) Immunoblots of HCT116 cells stably expressing the indicated variant of siRNA resistant FLAG-HA tagged HAPSTR1 and transfected with a control or HAPSTR1 targeting siRNA. Whole cell extracts were separated by SDS-PAGE and immunoblotted (IB) with the indicated antibodies. See also Figure S7.

To examine if nuclear HUWE1 was responsible for repressing p53 activity, we generated HCT 116 (wild type p53) cells with stable expression of siRNA resistant wild type, Δ1-102, or ΔNLS HAPSTR1. The increase in p53 levels upon HAPSTR1 knockdown was rescued upon reexpression of wild type, but not Δ1-102 or ΔNLS, HAPSTR1 (Figure 7D and S7D). This result is consistent with the hypothesis that nuclear HUWE1 represses p53 levels. Indeed, loss of HUWE1 resulted in a further repression of cell proliferation upon cisplatin treatment in a p53-dependent manner (Figure S7E). Overall, our results demonstrate that HAPSTR1 acts to localize HUWE1 to the nucleus to regulate a likely broad swath of nuclear targets impacting inflammatory and p53 signaling pathways.

## Discussion

### HUWE1 substrate plasticity and pleiotropic functions

HUWE1 is an enigmatic ubiquitin ligase that has been implicated to regulate numerous cellular pathways including DNA repair, transcription, proliferation, apoptosis, and cell signaling (Qi et al., 2022). Further, HUWE1 has been described to both promote and restrict tumorigenesis, as well as play critical roles in neurodevelopmental pathways (Adhikary et al., 2005; Forget et al., 2014; Giles and Grill, 2020; Gong et al., 2020; Hao et al., 2012; Inoue et al., 2013; King et al., 2016; Kunz et al., 2020; Myant et al., 2017; Opperman et al., 2017; Peter et al., 2014; Yang et al., 2018; Zhao et al., 2009; Zhao et al., 2008). These pleiotropic functions for HUWE1 suggest that it targets a diverse, and likely context-specific set of substrates for ubiquitylation. Our unbiased proteomic approach demonstrating a surprising lack of common HUWE1 substrates supports the hypothesis that HUWE1 targets diverse substrates for degradation in a cell typespecific manner. It is also likely that a subset of other ligases acts on a similarly broad set of substrates which may compensate for loss of HUWE1 in our and other studies. Indeed, our demonstration that overexpression of catalytically inactive HUWE1 results in a greater accumulation of substrates compared to the accumulation observed following reduction of HUWE1 protein levels, supports the idea that HUWE1 acts in concert with other ligases. Employing our proteomic pipeline with other proposed broad-acting ubiquitin ligases and examining substrate accumulation when combined with HUWE1 depletion will begin to address redundancy between pleiotropic ubiquitin ligases that display substrate selection plasticity.

### HUWE1 substrate targeting mechanisms

How broadly acting ubiquitin ligases target their substrates is an open question. Despite amazing progress in our understanding of protein ubiquitylation and degradation mechanisms, the substrate targeting mechanisms of only a small fraction of the over 600 ubiquitin ligases encoded within the human genome have been studied in detail. Previous structural studies of full-length HUWE1 revealed a surprising ring structure with the catalytic HECT domain positioned above the plane of the ring (Grabarczyk et al., 2021; Hunkeler et al., 2021). The diverse domain organization within HUWE1 includes three different ubiquitin-binding domains, a WWE domain, and a BH3 domain. It is suggested that HUWE1 recruits diverse substrates using these domains. The demonstration that the HUWE1 armadillo repeats can engage the flexible N-terminus of DDIT4 indicates that HUWE1 can also bind substrates within the central cavity of the structural ring (Hunkeler et al., 2021). These observations establish a possible mechanism in which HUWE1 engages a structurally highly diverse set of substrates and positions them within the central cavity to allow the HECT domain, which is positioned above the ring, to catalyze ubiquitylation.

The budding yeast HUWE1 ortholog, Tom1, also contains a ubiquitin binding domain and armadillo repeats, suggesting that the overall structure and substrate targeting mechanisms may be conserved. The expansion of the ubiquitin binding domains within human HUWE1 suggest that HUWE1 may engage substrates that were previously ubiquitylated by other ubiquitin ligases. Tom1 was recently shown to act in an E4 like mechanism to target a model ubiquitin-fusion degradation (UFD) substrate for degradation (Kats et al., 2022). In addition, HUWE1 and TRIP12, another pleotropic HECT-domain ligase, were demonstrated to target a UFD reporter substrate in human cells, arguing that HUWE1 may amplify ubiquitin chains on substrates previously ubiquitylated by other ligases (Poulsen et al., 2012). Binding preexisting ubiquitin chains may also allow HUWE1 to build branched ubiquitin chains on substrates to accelerate proteasome targeting, a property that has been described for HUWE1, TRIP12 and the HUWE1 ortholog Tom1 (Kaiho-Soma et al., 2021; Kats *et al.*, 2022; Kolla et al., 2022; Ohtake *et al.*, 2016; Yau et al., 2017). This possible E4 mechanism for HUWE1 is also consistent with the notion that HUWE1 may not have many dedicated substrates.

### HAPSTR1 regulates HUWE1 nuclear activity

The positive genetic relationship between HUWE1 and HAPSTR1 suggested that HAPSTR1 regulates HUWE1 function. Our findings that HAPSTR1 acts to localize HUWE1 to the nucleus and that nuclear HUWE1 is required for cell proliferation explains the genetic relationship. Our finding that HAPSTR1 is both a HUWE1 substrate and a HUWE1 regulator suggests a negative feedback mechanism in which HAPSTR1 recruits HUWE1 to the nucleus to target diverse nuclear substrates, including HAPSTR1 itself. Investigating how HAPSTR1 levels vary across cells and tissues would highlight contexts in which nuclear HUWE1 activity may be consequential for cell or tissue function.

Loss of HUWE1 or HAPSTR1 impacts diverse transcriptional pathways, including NF-κB-mediated inflammatory signaling. The mechanism by which HUWE1 activates baseline inflammatory signaling remains largely uncharacterized. While loss of HUWE1 and HAPSTR1 represses the transcription of many chemokines and cytokines, HAPSTR1 depletion was also shown to reduce overall chemokine secretion and cell migration (Amici et al., 2022). Our analysis also revealed enrichment for cell motility factors in transcripts repressed by loss of HAPSTR1 which suggests that cell migration may be broadly impacted by transcriptional programs regulated by nuclear HUWE1 activity. HUWE1 mutations have been linked to both syndromic and non-syndromic forms of human intellectual disability (ID) disorders (Froyen et al., 2012; Froyen et al., 2008; Giles and Grill, 2020; Madrigal et al., 2007; Opperman et al., 2017). Disease-linked mutations occur throughout HUWE1, including the ubiquitin-binding domains, but cluster within the HECT domain. These mutations largely impair HUWE1 activity in vitro and fail to rescue neuronal defects in *eel-1 (C.* elegans HUWE1 ortholog) deficient worms (Hunkeler et al., 2021; Opperman et al., 2017). As neural migration defects during development have been implicated in a wide range of neurodevelopmental disorders, including ID (Ossola and Kalebic, 2021), it is possible that disease-associated mutations may also impair HUWE1 nuclear localization and its ability to regulate a cell motility transcriptional program

## Acknowledgments

This work was supported by an American Cancer Society postdoctoral fellowship (PF-19-072-01 – TBE) to J.K.M. and NIH grants R01GM127681 (E.J.B) and R01CA262188 (E.S.F). This work was supported by The G. Harold and Leila Y. Mathers Charitable Foundation (E.S.F) and a Chleck Foundation Fellowship (M.W.M.).

## Author Contributions

Conceptualization, J.K.M., E.S.F., and E.J.B.; Formal analysis, K.A.D, E.S.F., and E.J.B.; Investigation, J.K.M., X.G., K.A.D., M.W.M., C.Y.J., M.L., and E.J.B.; Resources, M.H.; Writing – Original Draft, E.J.B.; Writing – Review and Editing, J.K.M., M.H., E.S.F., and E.J.B.; Visualization, J.K.M. and E.J.B.; Supervision and funding acquisition, E.S.F. and E.J.B.

## STAR Methods

### Reagents

Chemical reagents were used at the following concentrations: Doxycycline, 5 ug/mL; Bafilomycin A1, 100 nM; MG132, 10 μM. TAK-243, 1 μM. Etoposide, 50 μM. Cisplatin, 12.5 μM.

### Plasmids

Plasmids containing doxycycline-inducible shRNAs were obtained from Dharmacon. cDNAs in pDONR plasmids were verified by sequencing and cloned into expression vectors by Gateway cloning. HUWE1 variants were generated as described in (Hunkeler *et al.*, 2021). To generate siRNA resistant HAPSTR1 cDNA, silent mutations were introduced within the region recognized by HAPSTR1 siRNA oligo #1 by site-directed mutagenesis. For transient transfections, constructs were cloned into pDEST_CMV_FLAG. For stable expression, constructs were cloned into pDEST_pHAGE_FLAG-HA or pDEST_pHAGE_GFP.

### Cell lines, transfections and siRNA

293T, CAL27, and HCT116 cells were grown in DMEM (high glucose, pyruvate and L-glutamine) supplemented with 10% fetal bovine serum, 100 U/mL penicillin and 100 U/mL streptomycin. HAP1 cells were grown in IMDM supplemented with 10% fetal bovine serum, 100 U/mL penicillin and 100 U/mL streptomycin. HAP1 HUWE1 KO clone3 and clone5 were gifts from David Toczyski (UCSF). NCI-H2052 cells were grown in RPMI supplemented with 10% fetal bovine serum, 2 mM glutamine, 100 U/mL penicillin and 100 U/mL streptomycin. RPE1 cells were grown in DMEM:F-12 supplemented with 10% fetal bovine serum, 100 U/mL penicillin and 100 U/mL streptomycin. All cells were cultured at 37 °C with 5% CO_2_ and tested for mycoplasma contamination monthly.

Cell lines with stable transgene expression were generated by lentiviral infection followed by puromycin (inducible shRNA and pHAGE_FLAG-HA plasmids) or blasticidin selection (pHAGE_GFP plasmids).

Lipofectamine 2000 (Thermo Fisher) was used according to the manufacturer’s protocol for transient transfections of plasmids or co-transfection of plasmid and siRNA. Lipofectamine RNAiMAX (Thermo Fisher) was used according to the manufacturer’s protocol for siRNA transfections.

### In vitro ubiquitylation assay

0.5 uM UBE2D2 (R&D Systems) was charged with ubiquitin for 30 minutes by the addition of 1X E3 Ligase Conjugation Buffer (R&D Systems), 0.2 uM UBE1 (R&D Systems), 50 uM ubiquitin (R&D Systems), and 10 mM MgATP (R&D systems). In vitro ubiquitylation reactions were initiated by addition of 2 uM Strep-C16orf72/HAPSTR and 0.2 uM HUWE1 and allowed to proceed at the indicated temperature for 20 minutes. Reactions were quenched by addition of reducing SDS dye and analyzed via immunoblot with a 1:16,000 dilution of an anti-Strep HRP conjugate (Fisher Scientific). Blots were imaged on an Amersham Imager 600 using Amersham ECL Prime Western Blotting Detection Reagent (GE Life Sciences).

### HAP1 HUWE1 C4341S knock-in clone generation

Hap1 HUWE1-C4341S-KI was generated via prime editing at the endogenous site. The pegRNA sequence was designed with an online tool (http://deepcrispr.info/DeepPE/) and the following sequences were selected: protospacer sequence: 5’-GCCTGCCTTCAGCTCACACA-3’; reverse transcription template including edit: 5’-CTTTTACgATGT-3’; primer-binding site: 5’-GTGAGCTGAAGGC-3’. The plasmid expressing pegRNA was cloned via Gibson assembly using a gBlock gene fragment containing the U6 promoter and the pegRNA sequence (purchased from IDT) and a linearized pBluescript vector. Hap1 cells were seeded in 6-well plates and transfected at roughly 70% confluency with lipofectamine 3000 (Invitrogen) according to the manufacturer’s instructions and 500 ng pegRNA plasmid and 1 μg PE plasmid expressing the prime editor (PE2-2A-GFP, Addgene #132776). Cells were cultured for 3 days following transfection and GFP+ single cells were isolated into individual wells of 96-well plates by fluorescence-activated cell sorting (FACS). Cells were expanded for 10 days before genotyping. Genomic DNA was extracted and the editing site was amplified with PCR using the forward primer 5’-GTCCCTTCCTACAGATCCAGTG-3’ and the reverse primer 5’-CATCAAGTATGCAAGCTCAACC-3’. The PCR products were first screened using digestion by NlaIII and the potential hits were verified by Sanger sequencing.

### Cal27-Cas9 clone, Cal27 iCas9 sgRNA and transgene-expressing cell line generation

Cal27 clones expressing doxycycline-inducible 3xFLAG-Cas9 were generated by lentiviral infection with pCW-Cas9-Blast (Addgene, #83481), selection with 5 ug/mL Blasticidin for seven days, and dilution to yield individual clones. Clones were analyzed for doxycycline-inducible Cas9 expression by western blot over several weeks. The iCas9 sgRNA pools were generated by lentiviral infection of the Cal27-Cas9 clone with LentiGuide-Puro expressing the specific gene-targeting Grna. The guide sequences were as follows: control guide 5’-GCATCGTACGCGTACGTGTT-3’, HUWE1 guide 5’-TCCAGTGCGAGTTATATCAC-3’, C16ORF72 guide 5’-CGGGAGGCCGCGGAGGATGG-3’. Plasmids expressing sgRNAs were constructed by ligation of the annealed oligonucleotides into linearized LentiGuide-Puro vector (Addgene, #52963). Integration of the sgRNA was selected with 1 μg/mL puromycin for seven days and the iCas9 sgRNA pool was analyzed for doxycycline-inducible Cas9 expression by western blot. The pool targeting HUWE1 was also analyzed for reduced Huwe1 expression by western blot. The transgene-expressing cell lines were generated with lentiviral infection of the iCas9 sgC16ORF72 pool with plasmids expressing the Flag-tagged transgene and mCherry fluorescence protein. The mCherry+ pools were isolated by FACS.

### GFP-based and internally controlled competitive growth assay

Constitutively GFP-expressing Cal27 cells were generated by lentiviral infection with GFP-IRES-Blast, selection with 5 ug/mL Blasticidin for seven days, and dilution to yield individual cells that highly expressed GFP. The clonal population was checked for sustained GFP expression by flow cytometry. The iCas9 sgRNA pools and GFP+ clone were first treated with 3 μg/mL doxycycline for 3 days individually. Then, the iCas9 sgRNA pools were mixed with the GFP+ clone at approximately equal GFP+ to GFP-ratios and the GFP-/GFP+ ratios were analyzed by flow cytometry on day zero when cells were mixed and day three. Data were analyzed using FlowJo and the GFP-/GFP+ ratios were normalized to the day zero ratio first. The ratios for sgRNA pools were further normalized to that of sgControl.

### Immunoblotting

Cell pellets were resuspended 8 M urea, 50 mM Tris pH 8.0, 75 mM NaCl, 1 mM β-glycerophosphate, 1 mM NaF, 1 mM NaV, 40 mM NEM and EDTA-free protease inhibitor cocktail (Roche Diagnostics), sonicated, and cleared by centrifugation at 15,000 rpm for 10 min at 4 °C. Protein concentrations were determined by BCA Protein assay (Thermo Fisher). Samples were separated by SDS-PAGE and transferred to PVDF membranes (Bio-Rad) with a semi-dry transfer apparatus (Bio-Rad Turbo Transfer). Membranes were blocked with 4% milk in TBST. Primary antibodies were diluted in 5% BSA in TBST. HRP-conjugated secondary antibodies were diluted in 4% milk in TBST. Immunoblots were developed with Clarity Western ECL Substrate (Bio-Rad) and imaged on a Bio-Rad Chemi-Doc XRS+ system.

### Nuclei isolation

Nuclei were isolated as described in (Luo et al., 2014). The following buffers were used: Low-salt wash buffer (10 mM HEPES-KOH pH 7.9, 10mM KCl, 1.5 mM MgCl_2_, 0.1 mM EDTA, 1 mM β-glycerophosphate, 1 mM NaF, 1 mM NaV, 40 mM NEM and EDTA-free protease inhibitor cocktail (Roche Diagnostics)), Hypo-osmotic lysis buffer (Low-salt wash buffer + 0.3 M sucrose, 2% (v/v) Tween 40), 1.5 M sucrose buffer (Low-salt wash buffer + 1.5 M sucrose), and High-salt extraction buffer (20 mM HEPES-KOH pH 7.9, 40 mM NaCl, 1.5 mM MgCl_2_, 0.2 mM EDTA, 25% glycerol). Buffers and samples were kept on ice throughout the protocol. Cell pellets were resuspended in hypo-osmotic lysis buffer and homogenized by pipetting 100 times using a micropipette with a 200-uL pipette tip. The samples were overlaid on 1 mL of 1.5 M sucrose buffer and centrifuged at 12,000 rpm for 10 minutes at 4°C. After centrifugation, a volume equal to the amount of lysis buffer used was pipetted off the top of sample and saved as the cytoplasmic fraction. The rest of the supernatant was discarded and the nuclear pellets were resuspended in 1 mL of low-salt wash buffer. The nuclei were pelleted by centrifugation at 12,000 rpm for 30 seconds at 4°C. The supernatant was removed and the washed nuclear pellets were resuspended in high-salt extraction buffer and incubated on ice for 20 minutes with occasional vortexing. The nuclei were centrifuged at 12,000 rpm for 20 minutes at 4°C. The supernatants were retained as high-purity nuclear proteins. Nuclear proteins were spotted onto PVDF using a dot blot apparatus. Cytoplasmic proteins were separated by SDS-PAGE.

### Immunofluorescence

Cells were grown on glass coverslips coated with poly-L-lysine and fixed with 4% formaldehyde in phosphate-buffered saline (PBS) for 10 min. Blocking and primary antibody dilutions were done in AbDil (20 mM Tris, 150 mM NaCl, 0.1% Triton X-100, 3% bovine serum albumin, and 0.1% NaN3, pH 7.5). PBS with 0.1% Triton X-100 (PBS-TX) was used for washes and secondary antibody dilutions. Hoechst-33342 was used to visualize DNA. Coverslips were mounted in Fluoromount-G and sealed with nail polish.

### Fluorescence microscopy

Images were acquired on a SP8 confocal microscope (Leica Microsystems) with a 63x objective. Images were maximally projected from five z-sections with 1 μm spacing. The fluorescence is scaled independently in each panel to better show the localization of each transgene.

DNA colocalization measurements were determined with CellProfiler image analysis software.

### Immunoprecipitations

For IP-MS experiments, FLAG-tagged HUWE1 or HAPSTR1 expressing cells were resuspended in 50 mM Tris pH 8.0, 200 mM NaCl, 2 mM TCEP, 0.1% NP-40, cOmplete™ Protease Inhibitor Cocktail (Roche Diagnostics). Cells were lysed by passing through a needle at least 20 times. Samples were centrifuged at 15,000 rpm for 10 min at 4 °C. Protein concentrations were determined by BCA Protein assay (Thermo Fisher). Equal concentrations of each sample were incubated with M2-FLAG magnetic beads for 1 hour at 4 °C. The beads were washed three times with 50 mM Tris pH 8.0, 2 mM TCEP, 0.1% NP-40, cOmplete™ Protease Inhibitor Cocktail (Roche Diagnostics). Protein was eluted by two incubations with occasional mixing in 0.1 M glycine HCl pH 2.7 for 20 minutes at room temperature. Elutions were treated with 35 mM TCEP for 30 minutes, followed by 150 mM iodoacetamide for 45 minutes, and then 200 mM DTT for 15 minutes. Proteins were precipitated with methanol/chloroform. Precipitated protein was resuspended in 200 mM EPPS pH 8.0. Proteins were digested with LysC and trypsin overnight at 37 °C. Formic acid was added to each sample to a final concentration of 0.7%. Samples were desalted using a SoLa plate (Thermo Scientific).

For IP samples that were analyzed by Western blotting, cells were lysed by incubating on ice in 50 mM Tris pH 8.0, 150 mM NaCl, 0.5% NP-40, 1 mM β-glycerophosphate, 1 mM NaF, 1 mM sodium orthovanadate, 40 mM NEM and cOmplete™ EDTA-free protease inhibitor cocktail (Roche Diagnostics). Lysates were cleared by centrifugation at 15,000 rpm for 10 min at 4 °C. Protein concentrations were determined by BCA Protein assay (Thermo Fisher). Equal concentrations of each sample were incubated with anti-HA agarose beads for 1 hour at 4 °C. The beads were washed four times with 50 mM Tris pH 8.0, 150 mM NaCl, 0.1% NP-40. To elute, beads were boiled in Laemmli sample buffer.

### Luciferase reporter assay

HEK293T cells were reverse transfected with siRNA using Lipofectamine RNAiMAX (Invitrogen) according to the manufacturer’s instructions and seeded in 24-well plates. 48 hours after siRNA transfection, cells were transfected with the reporter plasmid hRluc-NF-κB-firefly (Addgene, #106979) using Lipofectamine 2000 according to the manufacturer’s instructions. Some cells were treated with TNF-α (100ng/ml) for 6 hours before being harvested. 24 hours after plasmid transfection, luciferase activity was measured with the Dual-Glo Luciferase Assay System (Promega) according to the manufacturer’s instructions. Firefly luciferase activity was normalized to that of Renilla luciferase.

### qRT-PCR analysis

Total RNA was extracted from Cal27 cells with TRIzol reagent (Invitrogen). 1.5 μg of total RNA was reverse transcribed with the oligo(dT)_20_ and Super-Script III First-Strand Synthesis System (Life Technologies). qRT-PCR was performed using an iTaq SYBR Green mix (Bio-Rad) and the CFX96 Real-Time PCR Detection System (Bio-Rad). The samples were run in biological triplicates and normalized to the internal control GAPDH. The primers were as follows: *CCL20* forward, 5’-AACCATGTGCTGTACCAAGAG-3’; *CCL20* reverse, 5’-CAGTCAAAGTTGCTTGCTTCTG-3’; *CXCL3* forward, 5’-GCAGGGAATTCACCTCAAGA-3’; *CXCL3* reverse, 5’-GTGTGGCTATGACTTCGGTT-3’; *GAPDH* forward, 5’-GGTGGTCTCCTCTGACTTCAACA-3’; *GAPDH* reverse, 5’-GTTGCTGTAGCCAAATTCGTTGT-3’.

### TMT mass spectrometry

Cell pellets were resuspended in 8 M Urea, 50 EPPS pH 8.5, 50 mM NaCl, 1 mM β-glycerophosphate, 1 mM NaF, 1 mM sodium orthovanadate, and EDTA-free protease inhibitor cocktail (Roche Diagnostics). Cells were lysed with 10 passes through a 21G and cleared by centrifugation at 15,000 rpm for 10 min at 4 °C. Protein concentrations were determined by BCA Protein assay (Thermo Fisher). Lysates were treated with 10 mM TCEP for 30 minutes, followed by 15 mM iodoacetamide for 45 minutes, and finally 10 mM DTT for 15 min. 200 μg of protein for each sample was precipitated using methanol/chloroform as previously described (Donovan et al., 2018). Precipitated protein was resuspended in 8 M Urea, 50 mM HEPES pH 7.4, followed by dilution to 1.85 M urea with the addition of 200 mM EPPS, pH 8. Proteins were digested with LysC overnight at room temperature. The next day, the digestions were diluted two-fold with 200 mM EPPS pH 8 followed by digestion with trypsin for 6 hours at 37 °C. Anhydrous acetonitrile (ACN) was added to each sample to a final concentration of 10% v/v. Tandem mass tag (TMT) reagents (Thermo Fisher Scientific) were dissolved in ACN and added to each sample for 90 minutes at room temperature. The reaction was quenched with hydroxylamine for 15 minutes at room temperature. Each channel was combined in a 1:1 ratio, desalted using the Stage-Tip method and analyzed by LC-MS for channel ratio comparison. Samples were then combined using the adjusted volumes determined in the channel ratio analysis and dried down in a speed vacuum. The combined sample was then resuspended in 1% formic before desalting with C18 (Sep-Pak, Waters).

Data were collected using an Orbitrap Fusion Lumos mass spectrometer (Thermo Fisher Scientific, San Jose, CA, USA) coupled with a Proxeon EASY-nLC 1200 LC lump (Thermo Fisher Scientific, San Jose, CA, USA). Peptides were separated on a 50 cm 75 μm inner diameter EasySpray ES803a microcapillary column (Thermo Fisher Scientific). Peptides were separated over a 190 min gradient of 6 - 27% acetonitrile in 1.0% formic acid with a flow rate of 300 nL/min.

Quantification was performed using a MS3-based TMT method as described previously (McAlister et al., 2014). The data were acquired using a mass range of m/z 340 – 1350, resolution 120,000, AGC target 5 x 105, maximum injection time 100 ms, dynamic exclusion of 120 seconds for the peptide measurements in the Orbitrap. Data dependent MS2 spectra were acquired in the ion trap with a normalized collision energy (NCE) set at 35%, AGC target set to 1.8 x 104 and a maximum injection time of 120 ms. MS3 scans were acquired in the Orbitrap with HCD collision energy set to 55%, AGC target set to 2 x 105, maximum injection time of 150 ms, resolution at 50,000 and with a maximum synchronous precursor selection (SPS) precursors set to 10.

### TMT quantitative LCMS data analysis and statistical analysis

Proteome Discoverer 2.2 (Thermo Fisher Scientific) was used for .RAW file processing and controlling peptide and protein level false discovery rates, assembling proteins from peptides, and protein quantification from peptides. The MS/MS spectra were searched against a Swissprot human database (January 2021) containing both the forward and reverse sequences. Searches were performed using a 20 ppm precursor mass tolerance, 0.6 Da fragment ion mass tolerance, tryptic peptides containing a maximum of two missed cleavages, static alkylation of cysteine (57.02146 Da), static TMT labelling of lysine residues and N-termini of peptides (229.16293 Da), and variable oxidation of methionine (15.99491 Da). TMT reporter ion intensities were measured using a 0.003 Da window around the theoretical m/z for each reporter ion in the MS3 scan. The peptide spectral matches with poor quality MS3 spectra were excluded from quantitation (summed signal-to-noise across channels <100 and precursor isolation specificity < 0.5), and the resulting data was filtered to only include proteins with a minimum of 2 unique peptides quantified. Reporter ion intensities were normalized and scaled using in-house scripts in the R framework (R Core Team, 2014). Statistical analysis was carried out using the limma package within the R framework (Ritchie et al., 2015).

### LFQ quantitative mass spectrometry

Cell pellets were resuspended 8 M urea, 50 mM Tris pH 8.0, 150 mM NaCl, 1 mM β-glycerophosphate, 1 mM NaF, 1 mM sodium orthovanadate, 40 mM NEM and EDTA-free protease inhibitor cocktail (Roche Diagnostics), sonicated, and cleared by centrifugation at 15,000 rpm for 10 min at 4 °C. Protein concentrations were determined by BCA Protein assay (Thermo Fisher). 30 ug of each sample was treated with 10 mM TCEP for 10 minutes, followed by 15 mM iodoacetamide for 10 minutes. Proteins were precipitated with methanol/chloroform. Precipitated protein was resuspended in 8 M Urea, 50 mM Tris pH 7.8, followed by dilution to 1.85 M urea with the addition of 50 mM Tris pH 7.8. Proteins were digested with LysC for 2 hours at 37 °C. Samples were diluted two-fold with 50 mM Tris pH 7.8, followed by digestion with trypsin overnight at 37 °C. Formic acid/acetonitrile was added to each sample to a final concentration of 1.4%. Samples were desalted using the Stage-Tip method.

Data were collected using a TimsTOF Pro2 (Bruker Daltonics, Bremen, Germany) coupled to a nanoElute LC pump (Bruker Daltonics, Bremen, Germany) via a CaptiveSpray nanoelectrospray source. Peptides were separated on a reversed-phase C18 column (25 cm x 75 μm ID, 1.6 μM, IonOpticks, Australia) containing an integrated captive spray emitter. Peptides were separated using a 50 min gradient of 2 - 30% buffer B (acetonitrile in 0.1% formic acid) with a flow rate of 250 nL/min and column temperature maintained at 50 °C.

DDA was performed in Parallel Accumulation-Serial Fragmentation (PASEF) mode to determine effective ion mobility windows for downstream diaPASEF data collection (Meier et al., 2020). The DDA PASEF parameters included: 100% duty cycle using accumulation and ramp times of 50 ms each, 1 TIMS-MS scan and 10 PASEF ramps per acquisition cycle. The TIMS-MS survey scan was acquired between 100 – 1700 *m/z* and 1/k0 of 0.7 - 1.3 V.s/cm^2^. Precursors with 1 – 5 charges were selected and those that reached an intensity threshold of 20,000 arbitrary units were actively excluded for 0.4 min. The quadrupole isolation width was set to 2 *m/z* for *m/z* <700 and 3 *m/z* for *m/z* >800, with the *m/z* between 700-800 *m/z* being interpolated linearly. The TIMS elution voltages were calibrated linearly with three points (Agilent ESI-L Tuning Mix Ions; 622, 922, 1,222 *m/z)* to determine the reduced ion mobility coefficients (1/K_0_). To perform diaPASEF, the precursor distribution in the DDA *m/z*-ion mobility plane was used to design an acquisition scheme for DIA data collection which included two windows in each 50 ms diaPASEF scan. Data was acquired using sixteen of these 25 Da precursor double window scans (creating 32 windows) which covered the diagonal scan line for doubly and triply charged precursors, with singly charged precursors able to be excluded by their position in the m/z-ion mobility plane. These precursor isolation windows were defined between 400 - 1200 *m/z* and 1/k0 of 0.7 - 1.3 V.s/cm^2^.

### LFQ quantitative LCMS data analysis and statistical analysis

The diaPASEF raw file processing and controlling peptide and protein level false discovery rates, assembling proteins from peptides, and protein quantification from peptides was performed using library free analysis in DIA-NN 1.8. Library free mode performs an in silico digestion of a given protein sequence database alongside deep learning-based predictions to extract the DIA precursor data into a collection of MS2 spectra. The search results are then used to generate a spectral library which is then employed for the targeted analysis of the DIA data searched against a Swissprot human database (January 2021). Database search criteria largely followed the default settings for DIA including: tryptic with two missed cleavages, carbomidomethylation of cysteine, and oxidation of methionine and precursor Q-value (FDR) cut-off of 0.01. Precursor quantification strategy was set to Robust LC (high accuracy) with RT-dependent cross run normalization. Proteins with missing values in any of the treatments and with poor quality data were excluded from further analysis (summed abundance across channels of <100 and mean number of precursors used for quantification <2). Protein abundances were scaled using in-house scripts in the R framework (R Development Core Team, 2014) and statistical analysis was carried out using the limma package within the R framework (Ritchie *et al.*, 2015).

### RNA sequencing

Total RNA was extracted using TRIzol (Invitrogen) according to the manufacturer’s protocol. After TapeStation analysis to verify sample quality, mRNA stranded libraries were prepared and sequenced on a NovaSeq S4 with PE100. Sequencing reads were aligned to the human genome (BSgenome.Hsapiens.UCSC.hg19, splicedAlignment=FALSE) and quantified at the level of genes (TxDb.Hsapiens.UCSC.hg19.knownGene Bioconductor package) using the QuasR (Gaidatzis et al., 2015) package with default parameters. Differentially expressed Genes were identified using the edgeR Bioconductor package (Robinson et al., 2010).

