## Supplementary Material for "HAPSTR1 localizes HUWE1 to the nucleus to limit stress signaling pathways"

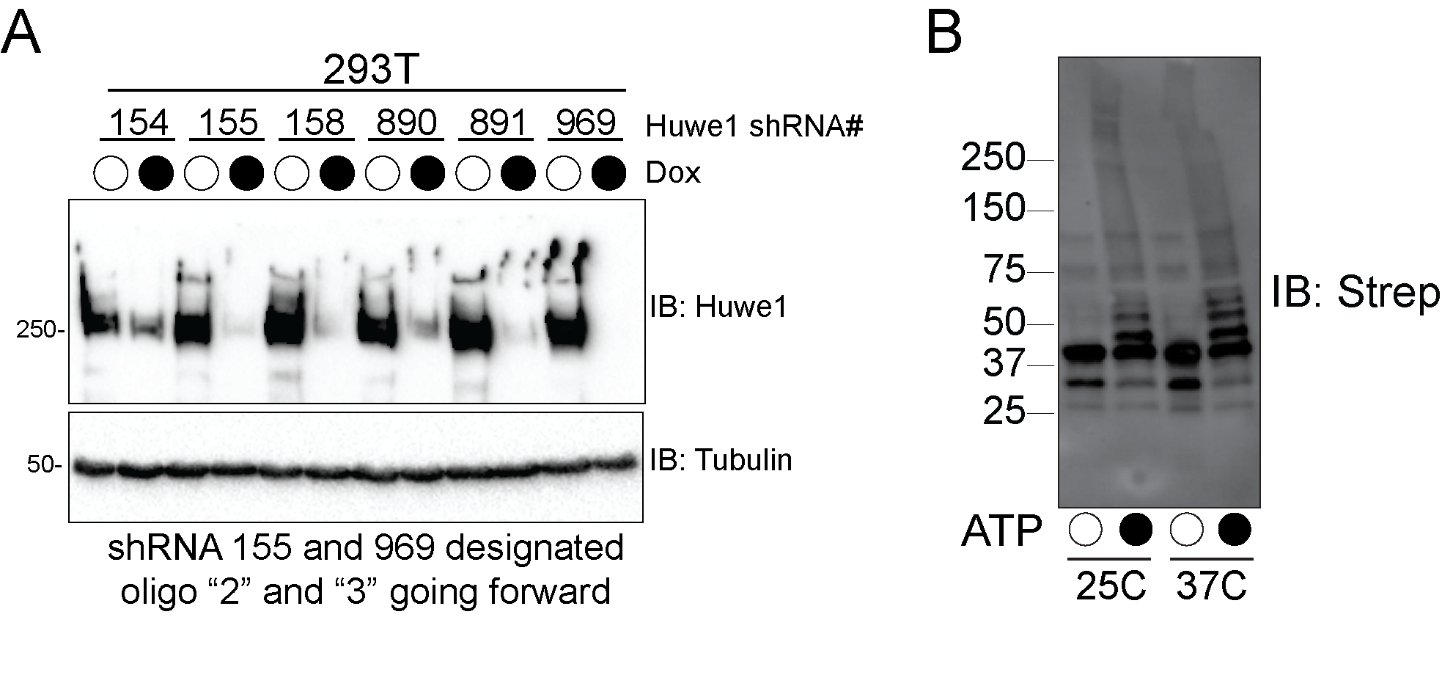

**Figure S1 – HUWE1 ubiquitylates HAPSTR1 *in vitro* (Related to Figure 1)**

A) Immunoblots of 293T cells stably expressing doxycycline-inducible shRNAs targeting HUWE1 were treated with doxycycline for 72hrs. Whole cell extracts were separated by SDS-PAGE and immunoblotted (IB) with the indicated antibodies.

B) Purified Strep-HAPSTR and HUWE1 were mixed with ubiquitylation components and either with and without ATP and incubated at the indicated temperature. Reactions were immunoblotted as indicated.

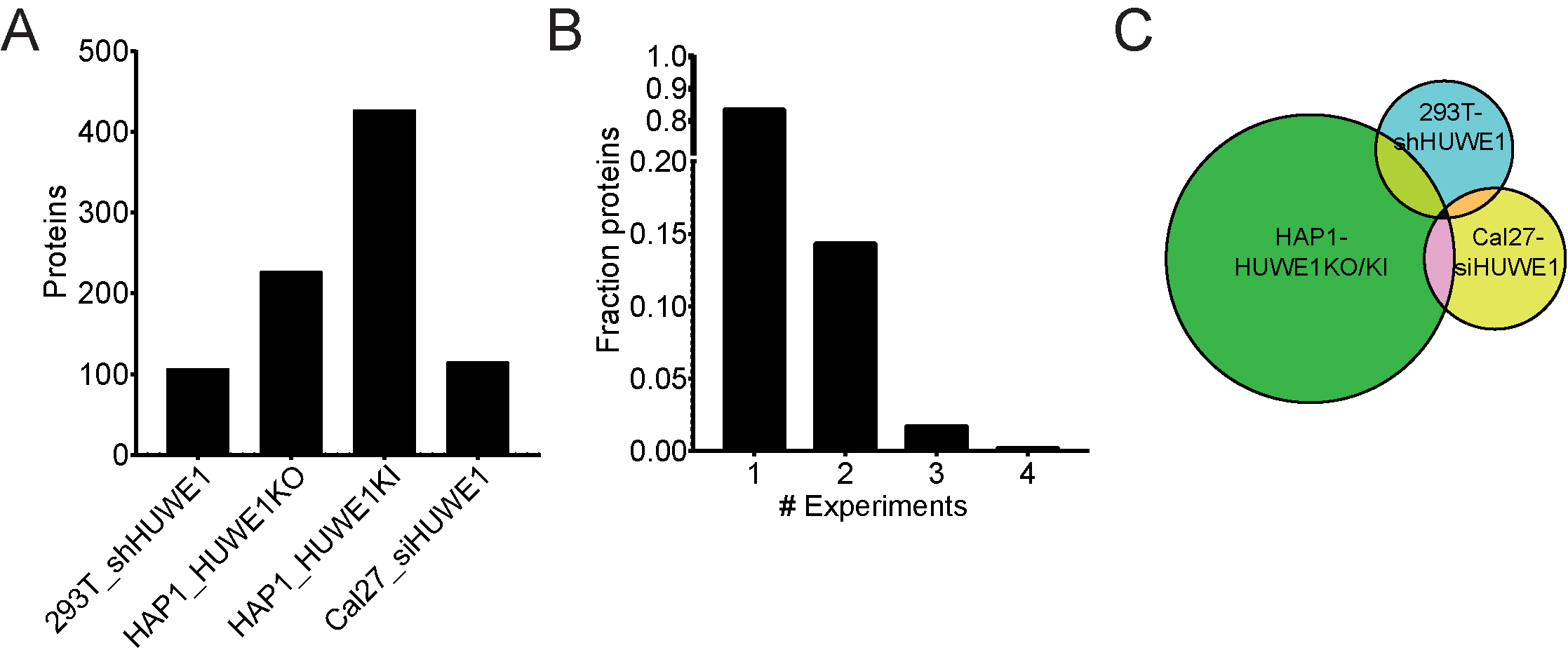

**Figure S2 – HUWE1 substrates are context-specific (Related to Figures 2 and 3)**

A) Number of proteins with significantly increased abundance in four different experiments with HUWE1 loss of function.

B) The fraction of all proteins that increased in abundance across different experiments depicted in A.

C) Venn diagram depicting the proteins with increased abundance in the quantitative proteomic analysis of 293T cells with inducible expression of shRNA targeting HUWE1, HAP1 cells with HUWE1 knocked out (KO) or with a mutation of the catalytic cysteine of HUWE1 knocked in (KI), or CAL27 cells transfected with siRNA targeting HUWE1.

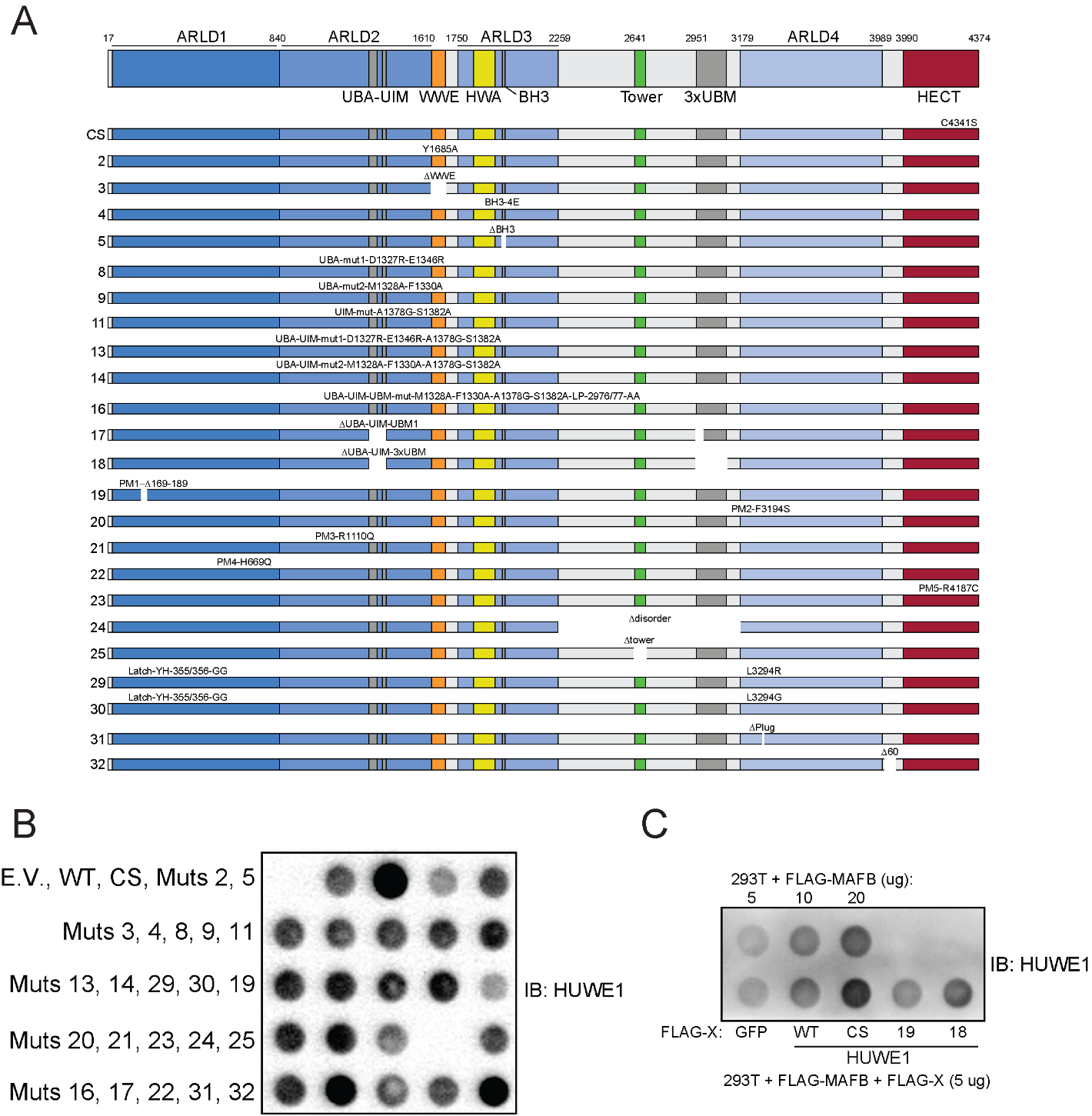

**Figure S3 – HUWE1 mutations differentially affect substrate degradation (Related to Figure 3)**

A) Schematic of HUWE1 depicting the domain organization and key structural features, and the mutations contained within each variant of HUWE1 used in this study. Variants 2-32 are referred to by their number throughout the manuscript.

B) Immunoblot of 293T cells transfected with an empty vector control (E.V.), wild type HUWE1 (WT) HUWE1 C4341S (CS, shown as mutant 1 in Fig. S3A) or the indicated HUWE1 variant as described in Fig. S3A. Whole cell extracts were spotted onto PVDF and immunoblotted with anti-HUWE1 antibody. The epitope recognized by the antibody is removed in Mutant 24.

C) Immunoblot of 293T cells transfected with FLAG tagged MAFB and either a FLAG tagged GFP control or the indicated HUWE1 variant. Whole cell extracts were spotted onto PVDF and immunoblotted for HUWE1.

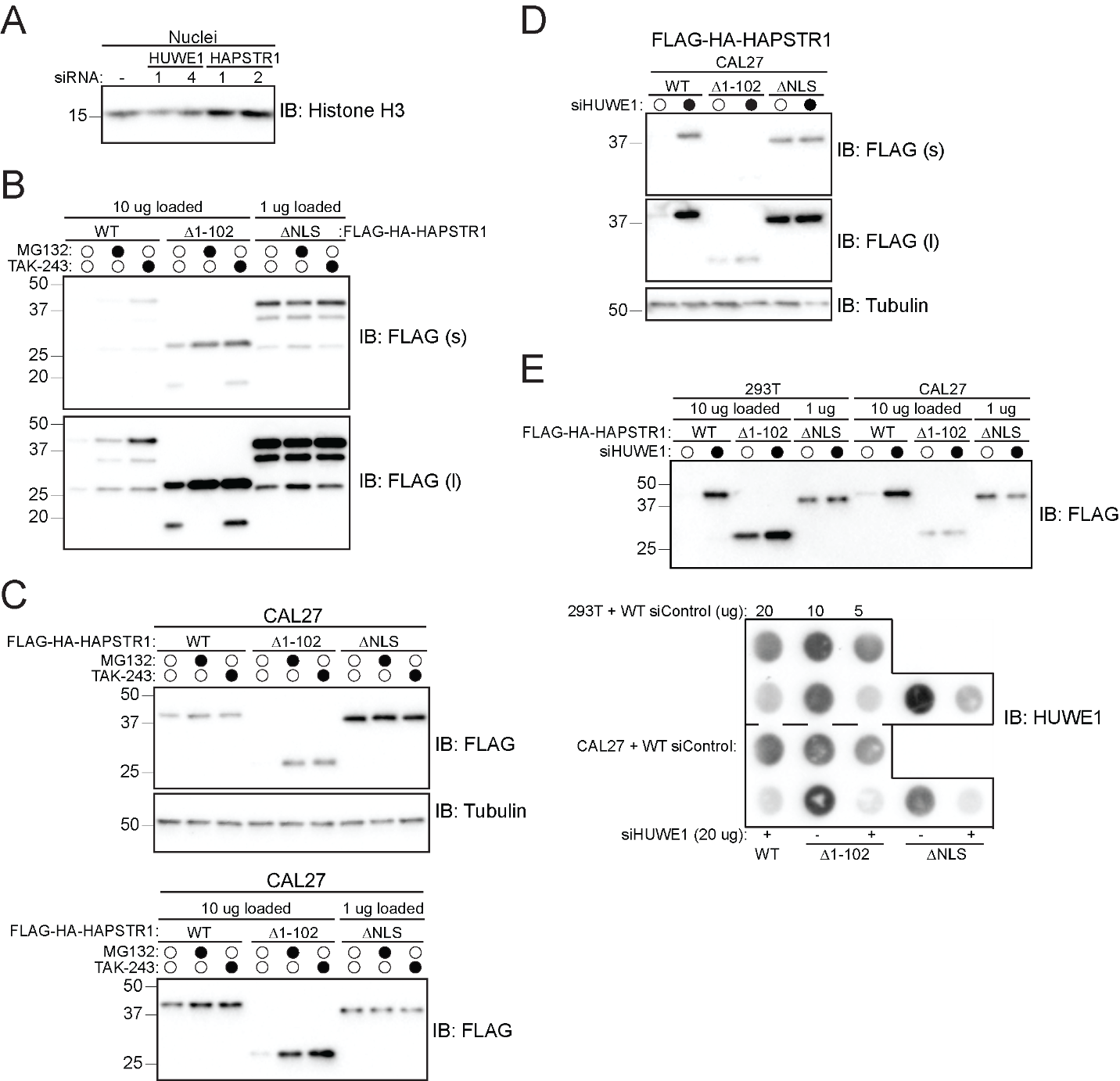

**Figure S4 – HAPSTR1 localizes HUWE1 to the nucleus (Related to Figure 4)**

A) Immunoblot of NCI-H2052 cells transfected with a control siRNA or one of two siRNA sequences targeting either HUWE1 or HAPSTR1 for 72 hours. The nuclei were isolated, separated by SDS-PAGE, and immunoblotted for histone H3.

B) Immunoblot of 293T cells stably expressing FLAG-HA tagged wildtype (WT) or the indicated variant of HAPSTR1 and treated with MG132 or TAK-243 for eight hours. Whole cell extracts were separated by SDS-PAGE and immunoblotted (IB) for FLAG (s, short exposure; l, long exposure).

C) Immunoblots of CAL27 cells stably expressing FLAG-HA tagged wildtype (WT) or the indicated variant of HAPSTR1 and treated with MG132 or TAK-243 for eight hours. Whole cell extracts were separated by SDS-PAGE and immunoblotted (IB) with the indicated antibodies. (Top) Equal amounts of protein were loaded for all samples. (Bottom) Ten-fold less protein was loaded for the ΔNLS samples relative to the wild type and Δ1-102 samples.

D) Immunoblots of CAL27 cells stably expressing FLAG-HA tagged wildtype (WT) or the indicated variant of HAPSTR1 (NLS, nuclear localization signal) and transfected with a control or HUWE1 targeting siRNA for 72 hours. Whole cell extracts were separated by SDS-PAGE and immunoblotted (IB) with the indicated antibodies (s, short exposure; l, long exposure).

E) Immunoblots of 293T or CAL27 cells stably expressing siRNA resistant FLAG-HA tagged wildtype (WT) or the indicated variant of HAPSTR1 (NLS, nuclear localization signal) were transfected with a control or HUWE1 targeting siRNA for 72 hours. (Top) Whole cell extracts were separated by SDS-PAGE and immunoblotted (IB) with the indicated antibodies. Ten-fold less protein was loaded for the ΔNLS samples relative to the wild type and Δ1-102 samples. (Bottom) Whole cell extracts were spotted onto PVDF and immunoblotted with anti-HUWE1 antibody.

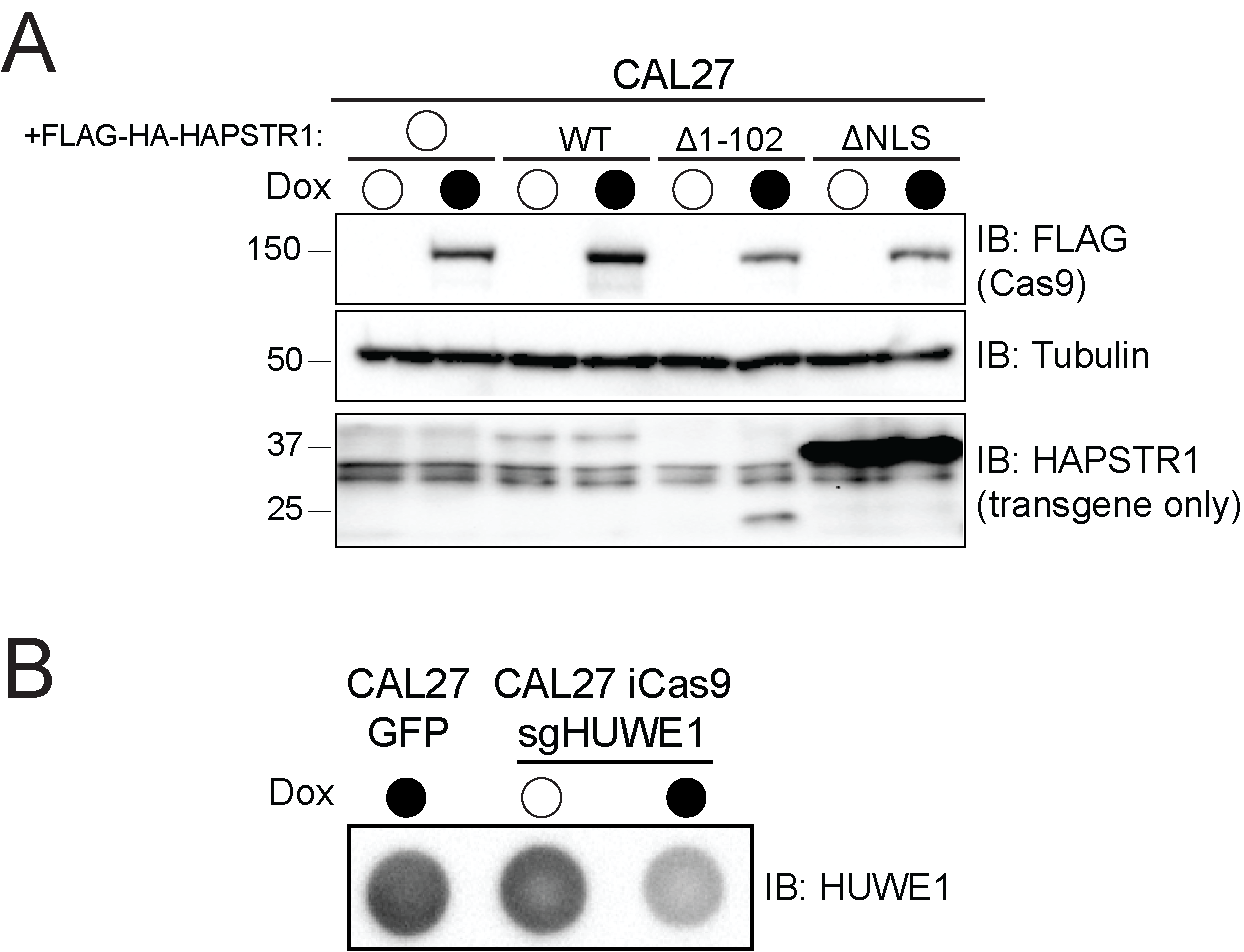

**Figure S5 – Nuclear HUWE1 is required for cellular proliferation (Related to Figure 5)**

A) Immunoblots of CAL27 cells stably expressing FLAG-HA tagged wildtype (WT) or the indicated variant of HAPSTR1 (NLS, nuclear localization signal), a sgRNA targeting HUWE1, and doxycycline-inducible Cas9 and treated with doxycycline for 72 hours. Whole cell extracts were separated by SDS-PAGE and immunoblotted (IB) with the indicated antibodies. The molecular weight of the band in the anti-FLAG blot corresponds to FLAG-Cas9 protein. FLAG-HAPSTR1 is significantly smaller and can be seen in the anti-HAPSTR1 blot. We note that the doublet bands are not endogenous HAPSTR1 as they are present even after knockdown of HAPSTR1 but RNAi (data not shown).

B) Immunoblot of CAL27 cells stable expressing GFP or CAL27 cells stably expressing a sgRNA targeting HUWE1 and doxycycline inducible Cas9 were treated with doxycycline for 72 hours. Whole cell extracts were spotted onto PVDF and immunoblotted with anti-HUWE1 antibody.

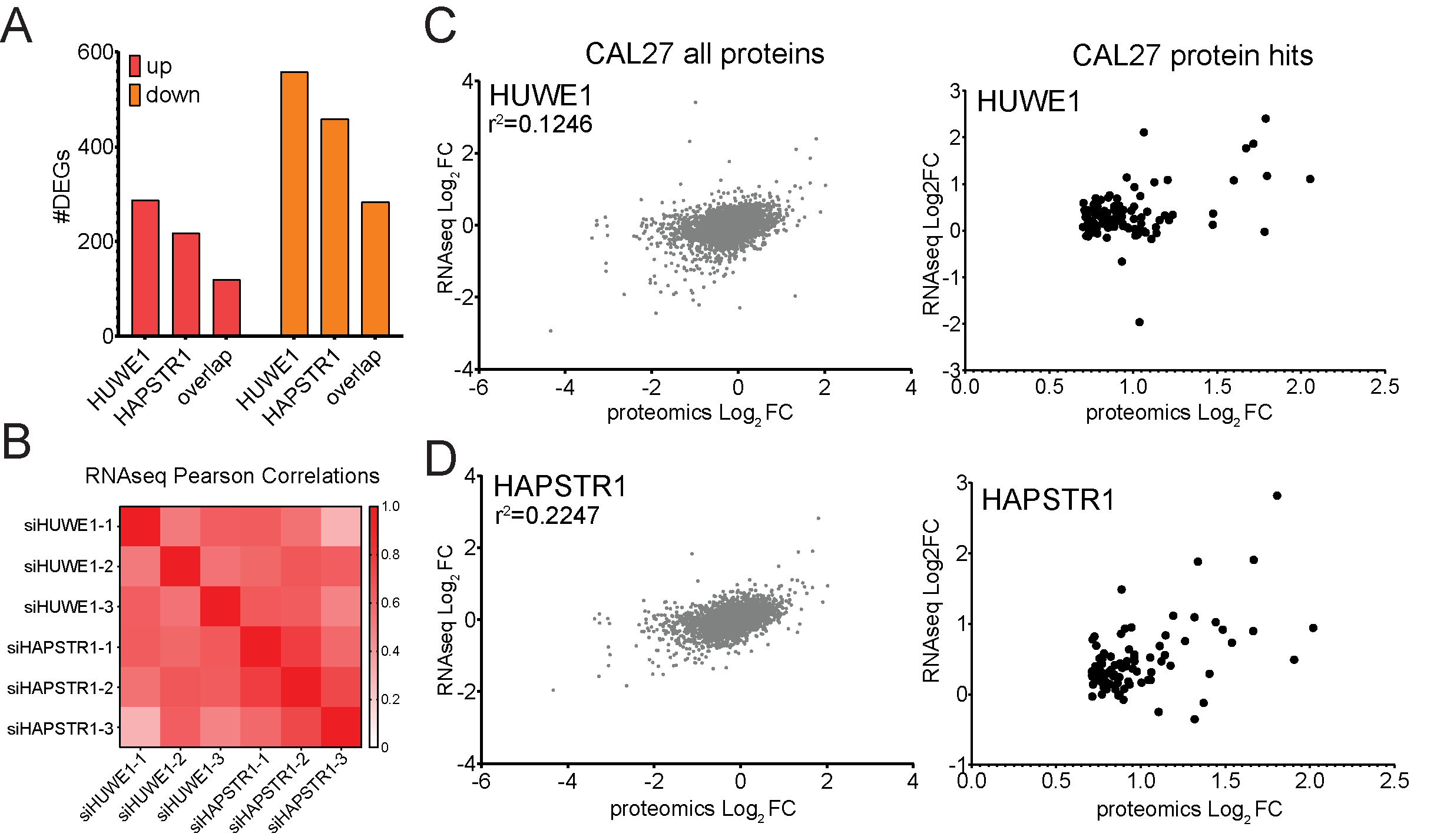

**Figure S6 – HUWE1 and HAPSTR1 regulate overlapping transcriptional programs (Related to Figure 6)**

A) CAL27 cells were transfected with a control siRNA or an siRNA targeting either HAPSTR1 or HUWE1 for 72 hours and mRNA was analyzed by RNA-seq. The number of differentially expressed genes in HUWE1 knockdown cells only, HAPSTR1 knockdown cells only, or in both HUWE1 and HAPSTR1 knockdown cells is shown. Genes with increased mRNA abundance are shown in red, genes with decreased mRNA abundance are shown in orange.

B) CAL27 cells were transfected with a control siRNA or an siRNA targeting either HAPSTR1 or HUWE1 for 72 hours and mRNA was analyzed by RNA-seq. The Pearson correlations comparing the differentially expressed genes across all pairwise combinations of HUWE1 and HAPSTR1 targeting siRNAs are shown.

C) Scatter plot comparing the log_2_ fold change (FC) for proteins quantified by label-free quantitative proteomics (x-axis) and the corresponding mRNA quantified by RNA-seq (y-axis) in CAL27 cells 72 hours after transfection of a HUWE1 targeting siRNA relative to a control siRNA. (Left) All proteins measured are shown. (Right) Only proteins with increased abundance after HUWE1 depletion are shown.

D) Scatter plot comparing the log_2_ fold change (FC) for all proteins quantified label-free quantitative proteomics (x-axis) and the corresponding mRNA quantified by RNA-seq (y-axis) in CAL27 cells 72 hours after transfection of a HAPSTR1 targeting siRNA relative to a control siRNA. (Left) All proteins measured are shown. (Right) Only proteins with increased abundance after HAPSTR1 depletion are shown.

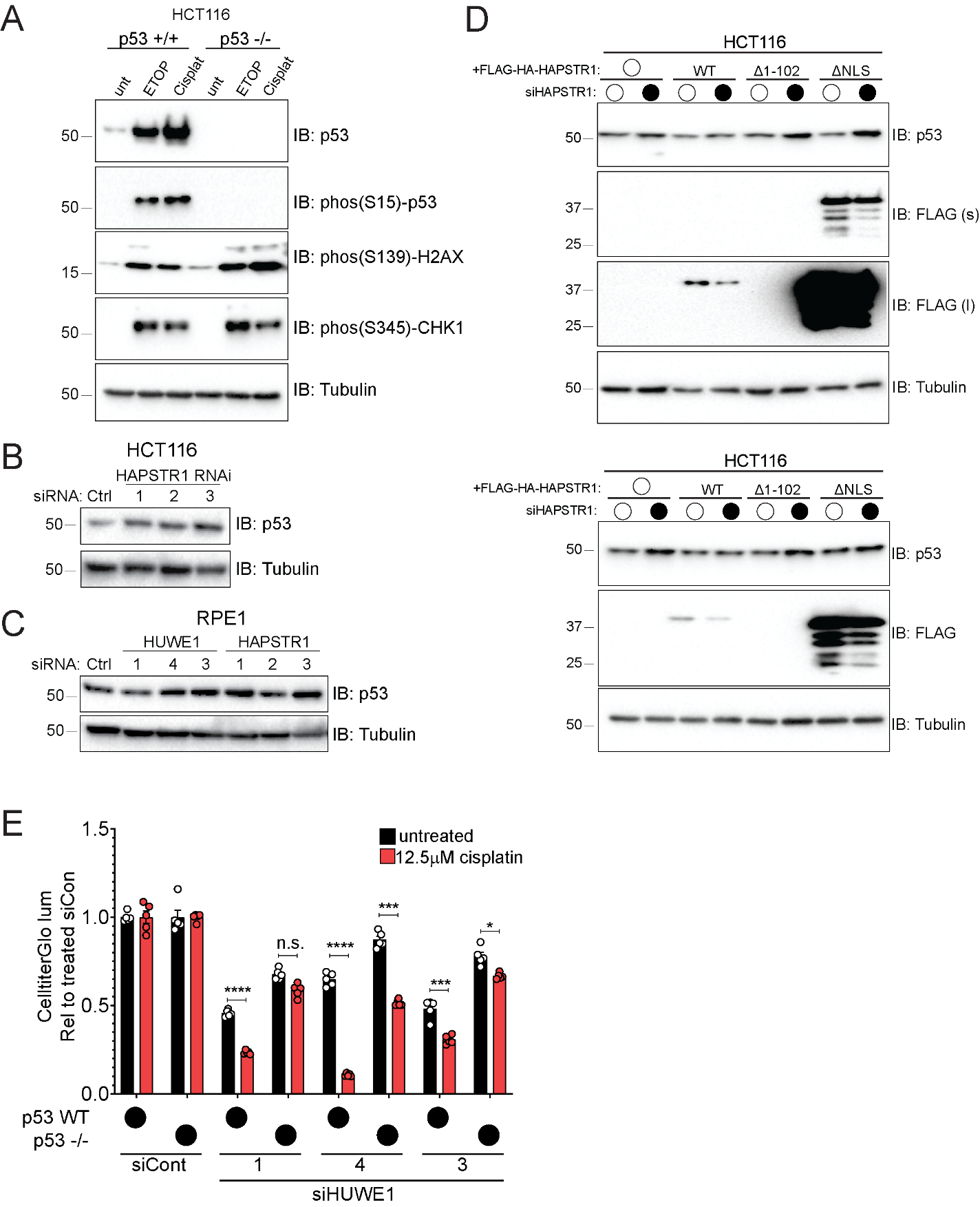

**Figure S7 – Nuclear HUWE1 regulates p53 (Related to Figure 7)**

A) Immunoblots of HCT116 cells with the indicated p53 status treated with etoposide (ETOP) or cisplatin (Cisplat). Whole cell extracts were separated by SDS-PAGE and immunoblotted (IB) with the indicated antibodies.

B) Immunoblots of HCT116 cells transfected with a control siRNA or one of three siRNA sequences targeting HAPSTR1. Whole cell extracts were separated by SDS-PAGE and immunoblotted (IB) with the indicated antibodies.

C) Immunoblots of RPE1 cells transfected with a control siRNA or one of three siRNA sequences targeting HUWE1 or HAPSTR1. Whole cell extracts were separated by SDS-PAGE and immunoblotted (IB) with the indicated antibodies.

D) Immunoblots of HCT116 cells stably expressing the indicated variant of siRNA resistant FLAG-HA tagged HAPSTR1 (WT, wild type; NLS, nuclear localization signal) and transfected with a control or HAPSTR1 targeting siRNA. Whole cell extracts were separated by SDS-PAGE and immunoblotted (IB) with the indicated antibodies. The blots represent separate replicate experiments. The p53 and tubulin blots from the top panel are duplicated from Figure 7D.

E) HCT116 cells with the indicated p53 status were transfected with a control (siCont) siRNA or one of three siRNA sequences targeting HUWE1. After 24 hours, cells were either untreated or treated with cisplatin and cell proliferation was measured 72 hours after drug treatment using the CellTiter-Glo assay. Mean values are shown with SEM. N=5. Statistical significance was determined by unpaired t-test comparing untreated and cisplatin treated conditions. **** p ≤ 0.0001; *** p ≤ 0.001; * p ≤ 0.05; n.s. p > 0.05.

| Mutant number ID | name | mutations | description |
| --- | --- | --- | --- |
| CS | C4341S | C4341S | Ligase inactive mutant |
| 2 | Y1685A | Y1685A | WWE domain mutation |
| 3 | delWWE | del1611-1683 | WWE domain deletion |
| 4 | BH3_4E | V1976E;L1980E;M1983E;V1987E | BH3 domain mutation |
| 5 | delBH3 | del1976-1990 | BH3 domain deletion |
| 8 | UBA_mut1 | D1327R;E1346R | UBA domain mutation |
| 9 | UBA_mut2 | M1328A;F1330A | UBA domain mutation |
| 11 | UIM_mut | A1378G;S1382A | UIM domain mutation |
| 13 | UBA-UIM_mut1 | D1327R;E1346R;A1378G;S1382A | UBA and UIM domain mutation |
| 14 | UBA-UIM_mut2 | M1328A;F1330A;A1378G;S1382A | UBA and UIM domain mutation |
| 16 | UBA-UIM-UBM_mut | M1328A;F1330A;A1378G;S1382A;L2976A;P2977A | UBA-UIM-UBM1-combined mutation |
| 17 | delUBA-UIM-UBM1 | del1318-1389;del2954-2989 | UBA-UIM-UBM1-combined deletion |
| 18 | delUBA-UIM-3xUBM | del1318-1389;del2954-3100 | UBA, UIM and all three UBM domain deletion |
| 19 | del169-189 | del169-189 | patient mutation 1 |
| 20 | F3194S | F3194S | patient mutation 2 |
| 21 | R1110Q | R1110Q | patient mutation 3 |
| 22 | H669Q | H669Q | patient mutation 4 |
| 23 | R4187C | R4187C | patient mutation 5 |
| 24 | deldisorder | del2260-3178 | large disorder domain deletion |
| 25 | delTower | del2641-2703 | deletion of structural tower |

**Table S4 – HUWE1 mutant IDs**
